# Content and performance of the MiniMUGA genotyping array, a new tool to improve rigor and reproducibility in mouse research

**DOI:** 10.1101/2020.03.12.989400

**Authors:** John Sebastian Sigmon, Matthew W Blanchard, Ralph S Baric, Timothy A Bell, Jennifer Brennan, Gudrun A Brockmann, A Wesley Burks, J Mauro Calabrese, Kathleen M Caron, Richard E Cheney, Dominic Ciavatta, Frank Conlon, David B Darr, James Faber, Craig Franklin, Timothy R Gershon, Lisa Gralinski, Bin Gu, Christiann H Gaines, Robert S Hagan, Ernest G Heimsath, Mark T Heise, Pablo Hock, Folami Ideraabdullah, J. Charles Jennette, Tal Kafri, Anwica Kashfeen, Samir Kelada, Mike Kulis, Vivek Kumar, Colton Linnertz, Alessandra Livraghi-Butrico, Kent Lloyd, Richard Loeser, Cathleen Lutz, Rachel M Lynch, Terry Magnuson, Glenn K Matsushima, Rachel McMullan, Darla Miller, Karen L Mohlke, Sheryl S Moy, Caroline Murphy, Maya Najarian, Lori O’Brien, Abraham A Palmer, Benjamin D Philpot, Scott Randell, Laura Reinholdt, Yuyu Ren, Steve Rockwood, Allison R Rogala, Avani Saraswatula, Christopher M Sasseti, Jonathan C Schisler, Sarah A Schoenrock, Ginger Shaw, John R Shorter, Clare M Smith, Celine L St. Pierre, Lisa M Tarantino, David W Threadgill, William Valdar, Barbara J Vilen, Keegan Wardwell, Jason K Whitmire, Lucy Williams, Mark Zylka, Martin T Ferris, Leonard McMillan, Fernando Pardo-Manuel de Villena

## Abstract

The laboratory mouse is the most widely used animal model for biomedical research, due in part to its well annotated genome, wealth of genetic resources and the ability to precisely manipulate its genome. Despite the importance of genetics for mouse research, genetic quality control (QC) is not standardized, in part due to the lack of cost effective, informative and robust platforms. Genotyping arrays are standard tools for mouse research and remain an attractive alternative even in the era of high-throughput whole genome sequencing. Here we describe the content and performance of a new Mouse Universal Genotyping Array (MUGA). MiniMUGA, an array-based genetic QC platform with over 11,000 probes. In addition to robust discrimination between most classical and wild-derived laboratory strains, MiniMUGA was designed to contain features not available in other platforms: 1) chromosomal sex determination, 2) discrimination between substrains from multiple commercial vendors, 3) diagnostic SNPs for popular laboratory strains, 4) detection of constructs used in genetically engineered mice, and 5) an easy to interpret report summarizing these results. In-depth annotation of all probes should facilitate custom analyses by individual researchers. To determine the performance of MiniMUGA we genotyped 6,899 samples from a wide variety of genetic backgrounds. The performance of MiniMUGA compares favorably with three previous iterations of the MUGA family of arrays both in discrimination capabilities and robustness. We have generated publicly available consensus genotypes for 241 inbred strains including classical, wild-derived and recombinant inbred lines. Here we also report the detection of a substantial number of XO and XXY individuals across a variety of sample types, the extension of the utility of reduced complexity crosses to genetic backgrounds other than C57BL/6, and the robust detection of 17 genetic constructs. There is preliminary but striking evidence that the array can be used to identify both partial sex chromosome duplication and mosaicism, and that diagnostic SNPs can be used to determine how long inbred mice have been bred independently from the main stock for a significant action of the genotyped inbred samples. We conclude that MiniMUGA is a valuable platform for genetic QC and important new tool to the increase rigor and reproducibility of mouse research.

## INTRODUCTION

The laboratory mouse is among the most popular and extensively used platforms for biomedical research. For example, in 2018 over 82,000 scientific manuscripts available in PubMed included the word “mouse” in the abstract. The laboratory mouse is such an attractive model due to the existence of hundreds of inbred strains and outbred lines designed to address specific questions, as well as the ability to edit the mouse genome; originally by homologous recombination and now with more efficient and simple techniques such as CRISPR (Dong *et al*. 2019; Ayabe *et al*. 2019). The centrality of genetics in mouse-enabled research begs the question of how genetic quality control (QC) is performed in these experiments.

We have a long track record of developing genotyping arrays for the laboratory mouse, from the Mouse Diversity Array (Yang *et al*. 2009) to the previous versions versions of the Mouse Universal Genotyping Array (MUGA,(Morgan *et al*. 2015)). These tools were originally designed for the genetic characterization of two popular genetic reference populations, the Collaborative Cross (CC) and the Diversity Outbred (DO), and then used for many other laboratory strains as well as wild mice (Yang *et al*. 2011; Collaborative Cross Consortium 2012; Carbonetto *et al*. 2014; Arends *et al*. 2016; Didion *et al*. 2016; Shorter *et al*. 2017; Srivastava *et al*. 2017; Rosshart *et al*. 2017; Veale *et al*. 2018). Efforts to extend the use of MUGA to characterize copy number variation and genetic constructs were met with limited success (Morgan *et al*. 2015). In conclusion, current genotyping tools are suboptimal for genetic QC and for new experimental designs aimed at facilitating the rapid identification of causal genetic variants in mouse crosses.

An improved genotyping platform would ideally be able to provide reliable information about the sex, genetic background and presence of genetic constructs in a given sample in a robust and cost-effective manner. The ability to discriminate between most genetic backgrounds is critical for genetic QC. The success of a new genotyping platform depends on how it compares to other more comprehensive solutions such as whole genome sequence (WGS) in terms of cost and ease involved in generating, analyzing, and interpreting the data. This is important because many analyses require more sophisticated approaches and skills that are beyond many users of laboratory mice. In addition, a new platform is needed to extend the success of reduced complexity crosses (RCC) beyond the C57BL/6J – C57BL/6NJ pair of strains (Kumar *et al*. 2013; Babbs *et al*. 2019). RCC are predicated on the idea that if a genetically driven phenotype is variable between a pair of closely related laboratory substrains, then QTL mapping combined with a complete catalog of the few thousand variants that differ among these substrains can lead to the rapid identification of the candidate causal variants (Kumar *et al*. 2013). This addresses one of the major limitations of standard mouse crosses, namely the cost in time and resources to move from QTL to quantitative trait variants (QTV). Genetic mapping in experimental F2 populations requires assigning every genomic region to one of three diplotypes based on their genotypes at segregating SNPs or other variants. The difficulty in RCC is two fold: first, genetic variants are unknown because WGS is not publicly available for most substrains; second, these variants are so rare (5-20K genome wide or one variant per 100 to 500 kb) that low pass WGS will miss the majority of them, complicating the analysis. In other words, the feature that makes RCC attractive for rapid QTV identification also makes it very difficult to implement.

To address these issues we created a fourth iteration of the MUGA family of arrays that we call MiniMUGA. The central considerations for the design were to reduce genotyping costs, provide broad discrimination between most inbred strains, support genetic mapping in dozens of different RCCs involving multiple substrains available from commercial vendors, robustly determine chromosomal sex, and reliably detect presence of popular genetic constructs. MiniMUGA fulfills all these criteria and facilitates simple, uniform and cost effective standard genetic QC, as well as serving the mouse community at large by providing a new tool for genetic studies.

## MATERIALS AND METHODS

### Reference samples

A diverse panel of 6,899 samples was used for calibrating and evaluating the performance of the array. The type of sample is provided in **Table 1**. To test the performance of each individual marker, provide reliable consensus genotypes and asses diagnostic markers, several biological and/or technical replicates for many inbred strains and F1 hybrids were included. **Supplementary Table 1** provides comprehensive information about each of these samples including sample name, type, whether it was genotyped in the initial or final version of the array, whether the sample was used to determine consensus genotypes or thresholds for chromosomal sex, chromosomal sex, basic QC metrics and values used to determine the presence of 17 constructs. A complete description of the information provided in **Supplementary Table 1** is available in the table legend.

**Table 1.**
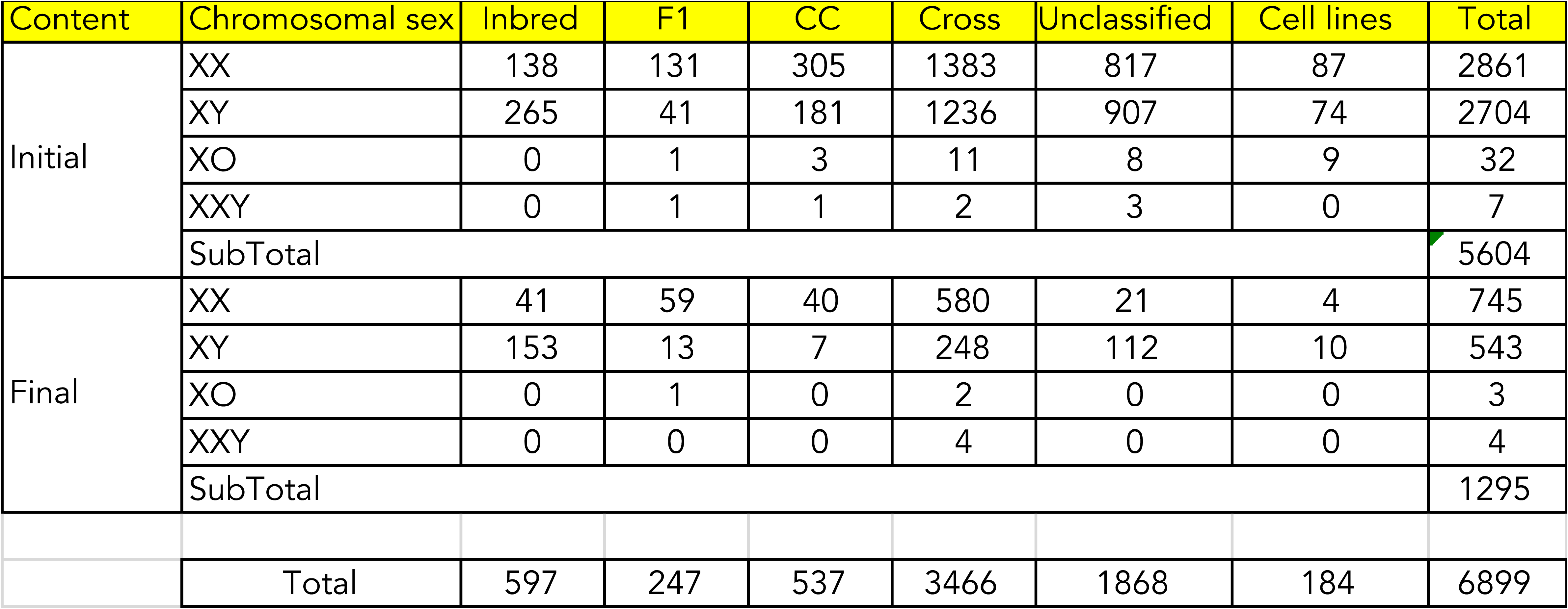

DNA stocks for classical inbred strains were purchased from The Jackson Laboratory or provided by the authors. DNA from most other samples was prepared from tail clips or spleens using the DNeasy Blood & Tissue Kit (catalog no. 69506; Qiagen, Hilden, Germany).

### Microarray platform

MiniMUGA is implemented on the Illumina Infinium HD platform (Steemers *et al*. 2006). Invariable oligonucleotide probes 50 bp in length are conjugated to silica beads that are then addressed to wells on a chip. Sample DNA is hybridized to the oligonucleotide probes and a single-basepair templated-extension reaction is performed with fluorescently labeled nucleotides. The relative signal intensity from alternate fluorophores at the target nucleotide is processed into a discrete genotype call (AA, AB, BB) using the Illumina BeadStudio software. Although the two-color Infinium readout is optimized for genotyping biallelic SNPs, both total and relative signal intensity can also be informative for copy-number variation and construct detection.

### Probe design

Of the 11,125 markers present in the array, 10,819 (97.2%) are probes designed for biallelic SNPs and the remaining 306 markers (2.6%) are probes designed to test the presence of genetic constructs (**Supplementary Table 2**). Nucleotides are labeled such that only one silica bead is required to genotype most SNPs, except the cases of [A/T] and [C/G] SNPs, which require two beads. In order to maximize information content, target SNPs were biased toward single-bead SNPs (mostly transitions). There are 10,721 single-bead assays and 404 two bead assays. The transition:transversion ratio in SNPs (excluding constructs) is 3:1.

### Array hybridization and genotype calling

Approximately 1.5 μg genomic DNA per sample was shipped to Neogen Inc. (Lincoln, NE) for array hybridization. Genotypes were called jointly for all reference samples using the GenCall algorithm implemented in the Illumina BeadStudio software.

### Probe Annotation

Probe design and performance of individual assays was used to annotate the array. **Supplementary Table 2** provides a rich set of annotations for each marker including: marker name, chromosome position, strand, probe sequence, performance, rsID, diagnostic value, thresholds for construct probes. A complete description of the information provided in **Supplementary Table 2** is available in the table legend.

### Chromosomal sex determination

We selected a set of 2,348 control samples (1,108 males and 1,240 females) with known X and Y chromosome number as determined through standard phenotypical sexing, which was supported by genotype analysis when expected heterozygosity on chromosome X was known. For each control sample, we first normalized the intensity values at each X and Y chromosome marker by dividing the intensity (r) by the sample’s median autosomal intensity. These autosome-normalized intensity values are used in all subsequent sex-determination calculations.

The first step of chromosomal sex determination was to identify sex-linked markers that provide a consistent estimate of of sex chromosome number with minimal noise. We identified 269 X and 72 Y sex-informative markers as those for which the ranges of median normalized intensity as defined by their standard deviations do not overlap between male and female controls (**Supplemental Figure 1**). This information is provided in **Supplementary Table 2**.

Next, we established chromosomal sex intensity threshold values. For each sample, we plotted the median of the normalized intensity values at the X informative markers on the x axis and median of the normalized intensity values at the Y informative markers on the y axis (**Figure 1**). Based on this plot we identified four clusters in sample intensity that correspond to XX, XY, XO, and XXY chromosomal sex. We defined thresholds as the midpoint between the relevant clusters. There is a single Y threshold value (0.3), separating samples with or without a Y chromosome. We identified two independent X threshold values (0.77 and 0.69) depending on whether the sample has a Y chromosome or not. These threshold values were used to classify the chromosomal sex of experimental samples into four groups, XX, XY, XO, or XXY.

**Figure 1.**
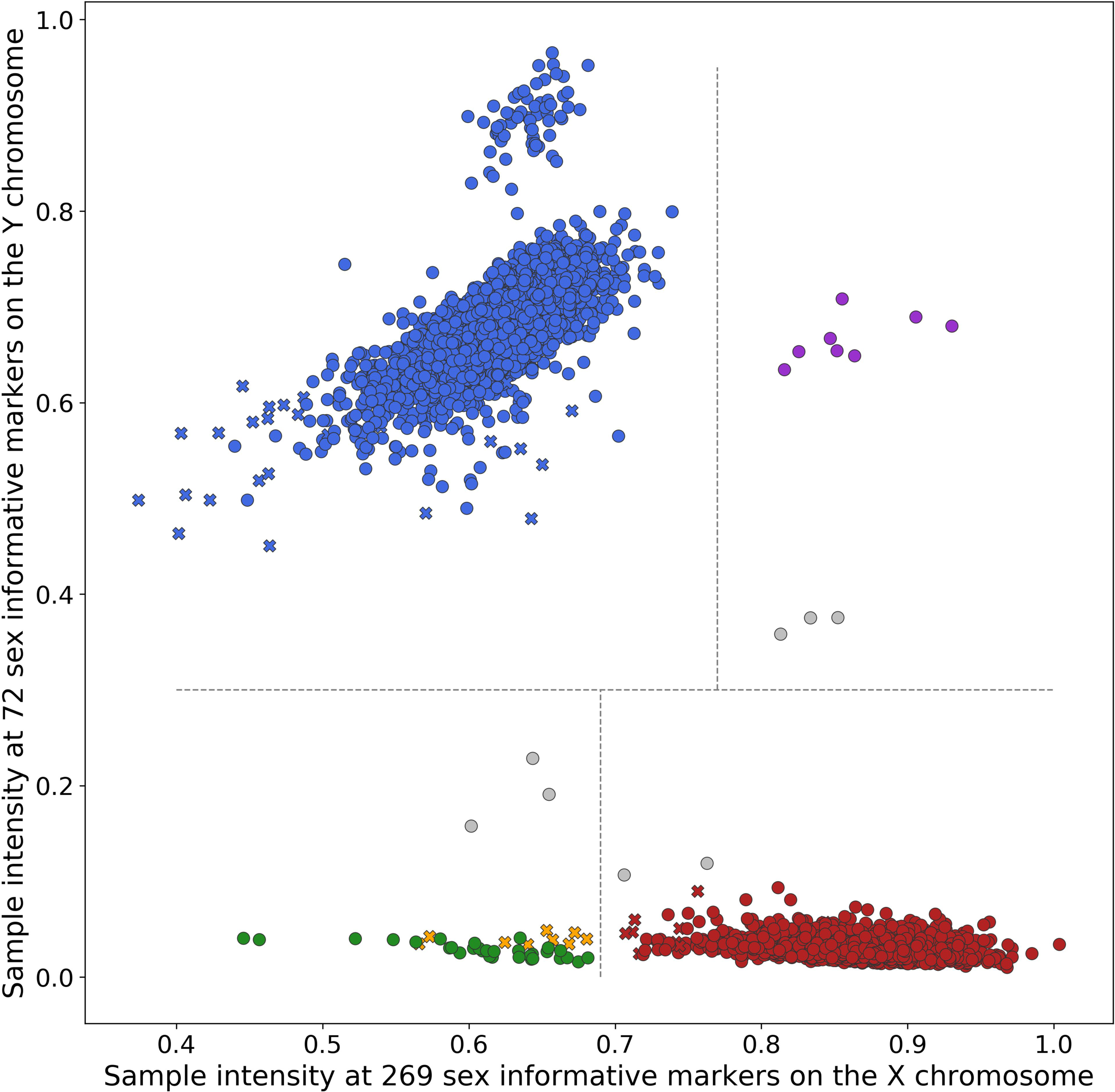
Chromosomal sex determination in 6,899 samples. Each dot and cross represents one sample. The x value is the autosome normalized median sample intensity at 269 sex informative X chromosome markers, and the y value is the autosome normalized median sample intensity at 72 sex informative Y chromosome markers. The dot color denotes the assigned chromosomal sex: XX, red; XY, blue; XO, green; XXY, purple. Potential mosaic samples are shown in gray and known errors in yellow. Samples with normal pd_stat as shown as circles and samples with high pd_stat are shown as crosses.

### Generation of consensus genotypes

The impetus for creating consensus genotypes for inbred strains in MiniMUGA is to provide a set of reference genotype calls for widely used strains. When possible, we included multiple biological and technical replicates of a given inbred strain to smooth over any errors in genotyping results, identify problematic markers, and to provide a more robust set of reference calls for comparison.

For each of 241 inbred strains (**Supplementary Table 3**), we genotyped from 1 to 19 samples (average 3.2 per strain). Most inbred strains (179) were genotyped more than once. For 53 strains (mostly BXD recombinant inbred lines) we did not genotype a male animal and thus Y chromosome genotypes are not provided for those strains. Over half of the strains (146) were genotyped only in the initial version of the array, so final content genotypes are missing in those strains. See **Supplementary Table 1** for details.

We generated consensus genotype calls at all 10,819 of the autosomal, X, pseudo-autosomal region (PAR), and Y chromosome markers (biallelic SNPs). For each consensus strain, at each marker, we recorded the genotype calls in all of the constituent samples and determined the consistency among these calls. For strains with more than one sample, if all calls are consistent, the consensus genotype is shown in upper case (A,T,C,G,H,N). Partially consistent calls are those with a mix of one or more calls of a single nucleotide and one or more H and/or N calls. Partially consistent calls are shown in lower case, as are calls for strains with a single constituent sample. Inconsistent calls are those for which two distinct nucleotides calls are observed. For standard markers, inconsistent genotypes within a strain are is shown as N in the consensus. For partially diagnostic SNPs the consensus call is the diagnostic allele shown in lower case. For CC strains, inconsistent consensus genotypes are shown as H, as these markers can be heterozygous in such samples. For mitochondria and Y chromosome markers, consensus calls follow the same rules except H calls are treated as N. **Supplementary Table 4** provides a list of rules for generating all possible consensus calls. **Supplementary Table 5** provides a listing of the consensus genotypes.

### Informative SNPs between closely related substrains

To increase the specificity of MiniMUGA as a tool for discriminating between closely related inbred strains, we used public data from several other studies providing genotype or whole-genome sequence information (Yang *et al*. 2009; Keane *et al*. 2011; Adams *et al*. 2015; Morgan *et al*. 2015). Most importantly, we included SNP variants that are segregating between substrains. These SNPs were identified by whole genome sequencing of 33 substrains performed as part of two ongoing collaborations (contributed by either MTF, RSB and MTH, or MTF and CMS; **Table 2**). Finally, we included 339 variants discriminating substrains of C57BL/10 (provided by AAP, YR and CSP). Some of the 5,171 GigaMUGA probes included to cover the genome uniformly in classical and wild-derived inbred strains were also informative for substrains.

**Table 2.**
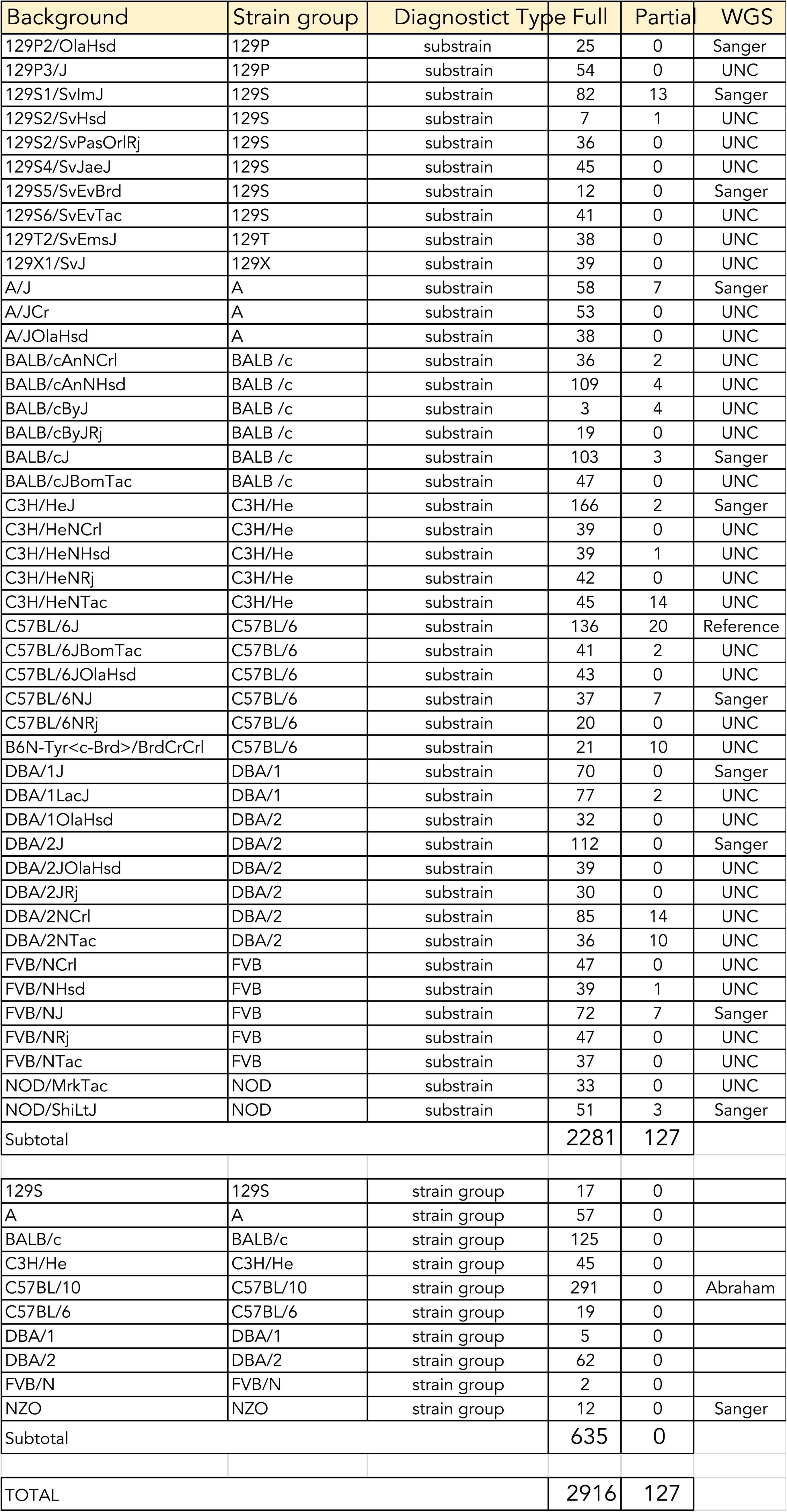

### Probes for genetically engineered constructs

We selected 36 constructs commonly used in genetic engineering in the mouse. For each construct, we obtained full length sequence from either Addgene or GenBank. We ran a BLAST search (Johnson *et al*. 2008) on these sequences to identify 2-5 additional sequences which (a) had high BLAST scores, and (b) were annotated as containing the relevant construct gene we were searching for (all sequence accession numbers are in **Supplementary Table 8**). For each construct, sequences were then aligned using the EMBOSS Water algorithm from EMBL-EBI (https://www.ebi.ac.uk/Tools/psa/emboss_water/). We identified conserved 50-mers within these alignments followed by a single A in the forward strand, or followed by a single T in the reverse strand. These sequences were submitted to the Illumina BeadStudio design pipeline, with a pseudo-SNP (A/G or T/C). Probes which passed a quality score threshold of 0.7 were included in the array. In total we created 306 probes for these constructs (range 3-18, median 8 probes/construct).

In order to validate these probes, we first eliminated probes which had high intensity signal in the 580 negative control samples (standard inbred mouse strains and F1 hybrids between them). Next, among the remaining probes, we identified those with significantly variable intensity among the remaining 6,319 samples in this study. In particular, we confirmed that, where available, positive controls had high signal intensity.

This process left 163 validated probes. We noticed that signal intensity of validated probes was often positively correlated with other validated probes with the same, or related target constructs. All validated probes were then subject to a second round of BLAST for final identification of the targeted constructs and to provide a biological basis for grouping of highly correlated probes. These alignments are provided in **Supplementary Figure 2.** In total these 163 probes mapped to 17 biologically distinct constructs (see **Table 3**). Probes tracking the hCMV enhancer can divided into two groups based on the clustering.

**Table 3.**
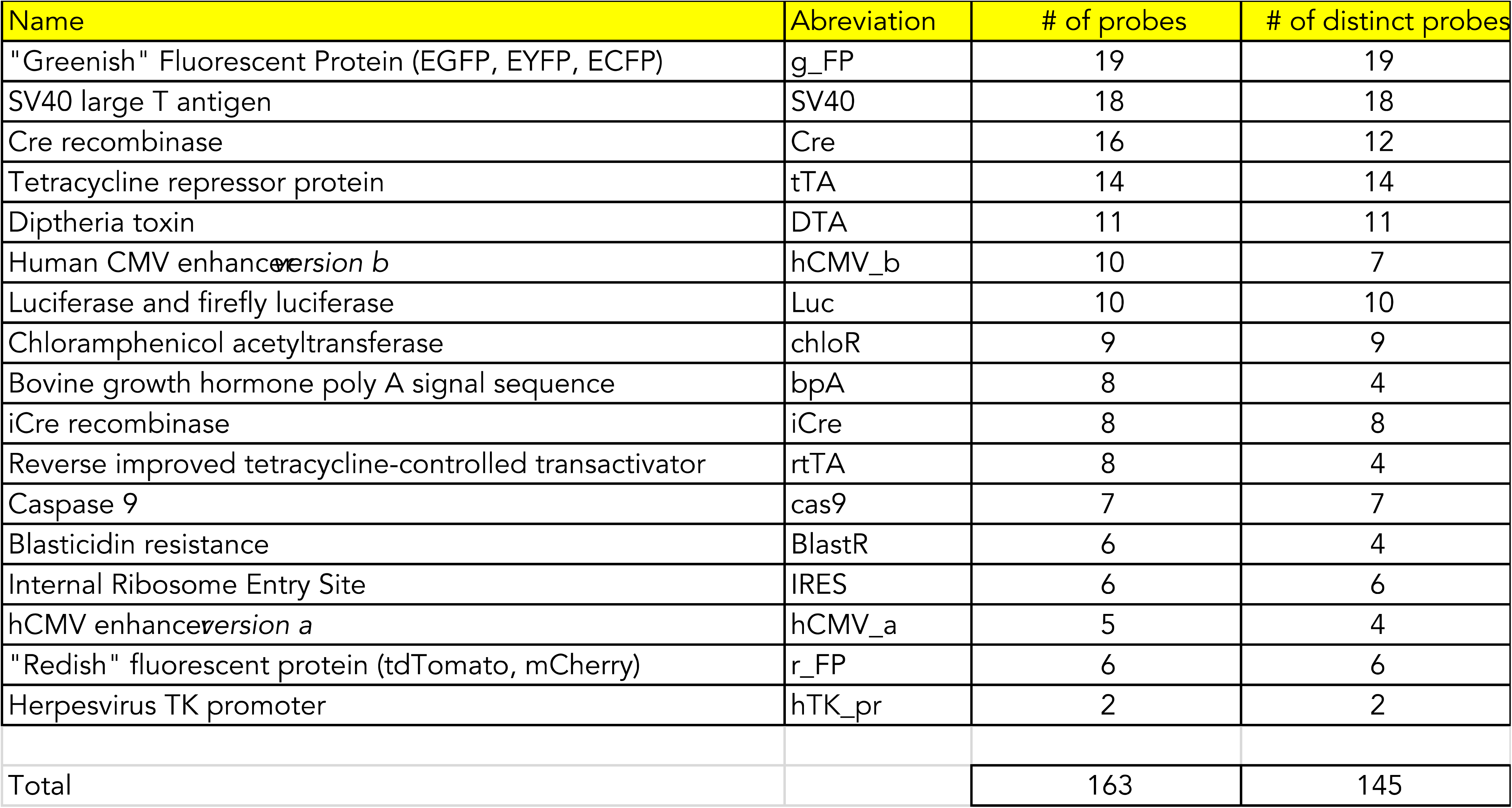

Once we selected the final set of validated probes for a specific construct, we used the per-sample distribution of the sum validated probe intensity to manually identify conservative threshold values for the presence and absence of each construct. We used the negative and positive controls to set initial thresholds and then used the distribution of values to identify breaks and set the final thresholds such that we minimize the number of samples misclassified as positive or negative.

### Additional sample quality metrics

Most quality metrics for genotyping arrays are based on genotype calls. However, intensity based analyses, such as chromosomal sex determination, assume quasi-normal distribution of marker intensities in a given sample (**Supplementary Figure 3**). In our dataset some samples had significantly skewed and idiosyncratic intensity distributions. Among these samples there is an excess of sex chromosome aneuploidies as called by our algorithm, many of which are in fact errors.

To identify samples with poor performance we first identified 200 random samples with no chromosomal abnormalities and confirmed that they have quasi-normal intensity distribution in aggregate. We then computed a Power Divergence statistic (pd_stat; equivalent to Pearson’s chi-squared goodness of fit statistic for each sample, comparing to that distribution. **Supplementary Figure 4** shows the distribution of pd_stat values in our entire dataset. We selected 3,230 as the threshold, such that in samples with higher values the reported chromosomal sex could be incorrect. This warning is particularly true for samples reported to have sex chromosome aneuploidy. The threshold also ensures that in samples from species other than *Mus musculus*, chromosomal sex determination is treated with skepticism.

To determine whether a high pd_stat had an effect on the accuracy of genotyping calls we selected four pairs of different F1 mice ((A/JxCAST/EiJ)F1_M15765; (CAST/EiJxA/J)F1_F002; (CAST/EiJxNZO/HlLtJ)F1_F0019; (CAST/EiJxNZO/HlLtJ)F1_F022; (NZO/HlLtJxNOD/ShiLtJ)F1_F0042; (NZO/HlLtJxNOD/ShiLtJ)F1_F0042; (PWK/PhJxNZO/HlLtJ)F1_F0019 and (PWK/PhJxNZO/HlLtJ)F1_M0001) that cover a variety of pd_stat comparisons (high/low, medium/medium, and low/low). For each pair we first determined the pairwise consistency of the genotypes calls and then compared these genotypes to predicted calls for the consensus reference inbred strains. Pairwise comparison consistencies in the autosomes excluding N calls vary between 99.5% and 100%. Similarly, the consistency with predicted genotypes is very high (99.5%-100%). We conclude that the pd_stat is independent of genotype call quality.

### Data availability

Genotype calls, hybridization intensity data and consensus genotypes for inbred strains (both raw and processed) for 6,899 samples are available for download at the Dataverse (upon acceptance flat files with the data will be posted).

## RESULTS

### Sample set, reproducibility and array annotation

To test the performance of the MiniMUGA array we genotyped 6,899 DNA samples from a wide range of genetic backgrounds, ages and tissues (**Supplementary Table 1**). These samples include many examples of inbred strains, F1 hybrids, experimental crosses and cell lines (**Table 1**). The array content was designed in two phases and thousands of samples were genotyped to determine the marker performance, information content and to improve different aspects of the proposed use of the array for genetic QC. In the initial array that contained 10,171 makers, 5,604 samples were genotyped. The second phase added 954 markers, with an additional 1,295 samples genotyped. This results in 6,300 samples that were genotyped once and 225 samples were genotyped two or more times, resulting in a total of 6,525 unique samples. The 599 replicates were used to estimate the reproducibility of the genotype data. Overall, 99.6 ± 0.4% of SNP genotype calls were consistent between technical replicates (range 95.9% to 100%). The consistency rate is similar for replicates run within and between versions of the array. Samples with lower consistency rates include wild-derived samples from more distant species and subspecies (SPRET/EiJ, SFM, SMZ, MSM/MsJ and JF1/Ms), lower quality samples, and cell lines. Inconsistency was typically driven by a small minority of markers and by “no calls” in one or few of the technical replicates.

Probe design and performance of individual assays was used to annotate the array. **Supplementary Table 2** contains the following information: 1) Marker name; 2) Chromosome; 3) Position; 4) Strand; 5-6) Sequences for one and two bead probes; 7-8) Reference and alternate allele at the SNP; 9) Tier; 10) rsID; 11) Diagnostic information; 12) Uniqueness; 13) X chromosome markers used to determine the presence and number of X chromosomes; 14) Y chromosomes markers used to determine the presence of a Y chromosome; 15) Markers added in the second phase.

### Improved chromosomal sex determination reveals sex chromosome aneuploidy due to strain-dependent paternal non-disjunction

Typically, genetic determination of sex of a mouse sample has relied on detecting the presence of a Y chromosome. This approach does not estimate X chromosome dosage and thus lacks the ability to identify samples with common types of sex chromosome aneuploidies. In contrast, MiniMUGA uses probe intensity to discriminate between normal chromosomal sexes (XX and XY) and two types of sex chromosome aneuploidies, XO and XXY (**Supplementary Table 1**). The methodology (Materials and Methods) relies on median autosome-normalized intensity at 269 X chromosome markers and 72 Y chromosome markers. This approach provides a robust framework to discriminate between at least four types of chromosomal sex (**Figure 1**). Our set of 6,899 samples was composed of 3,507 unique females (no Y chromosome present) and 3,018 unique males (Y chromosome present).

We initially identified 54 samples as potential XO and XXY. However, in eight XO females the pattern of heterozygosity and recombination in the X chromosome (**Supplementary Table 6**) demonstrates that these are, in fact, normal XX females with abnormal intensities. We developed a new QC test (pd_stat, see Materials and Methods) to identify samples in which chromosomal sex determination is not accurate. Once these eight samples were removed, 46 samples that had sex chromosome aneuploidies remained. To determine the rate of aneuploidy we only considered unique samples (not replicates). This results in 45 aneuploid samples among 6,525 total unique samples, an overall 0.7% rate. This rate is driven by a highly significant excess (7X) of sex chromosome aneuploids among the cell lines. Notably all these aneuploids are XO. Among live mice there were 36 unique aneuploids and the rate is 0.55%, similar but higher than the reported rate in wild mice and in humans (Searle and Jones 2002; Chesler *et al*. 2016; Le Gall *et al*. 2017). In this dataset, XO females are observed at significantly higher frequency than XXY males (p=0.02; 25 XO females and 11 XXY males) (**Table 1**).

For 22 of the 45 unique samples with sex chromosome aneuploidies, both parents were known and informative for the X chromosome. This information allowed us to potentially determine the parental origin of the missing (in XO) or the extra (in XXY) X chromosome based on the haplotype inherited and recombination patterns observed (**Supplementary Table 6; Figure 2**). Overall, the parental origin can be determined unambiguously in 21 of these samples, and in all but one sample (95%) the aneuploidy is due to sex chromosome non-disjunction in the paternal germ line (**Figure 2**). Note that this applies to both XO and XXY samples. Given the paternal origin of most sex chromosome aneuploidies, we investigated whether the type of sire had an effect. We observed a highly significantly (p<0.00001) excess of aneuploids in the progeny of (CC029/Unc x CC030/GeniUnc)F1 hybrid males than in all other sires. Out of 180 male progeny of this cross, 5% of genotyped samples were aneuploids and both XO and XXY were observed (3 XO and 6 XXY mice, respectively). There was also evidence of an excess of sex chromosome aneuploids in progeny of sires with CC011/Unc background (5 XO females, **Supplementary Table 6**). We conclude that sex chromosome aneuploidy is relatively common in lab mice, originates predominantly in the paternal germ line and depends on the sire genotype. In some backgrounds aneuploidy rate is a factor of magnitude higher than in the general population.

**Figure 2.**
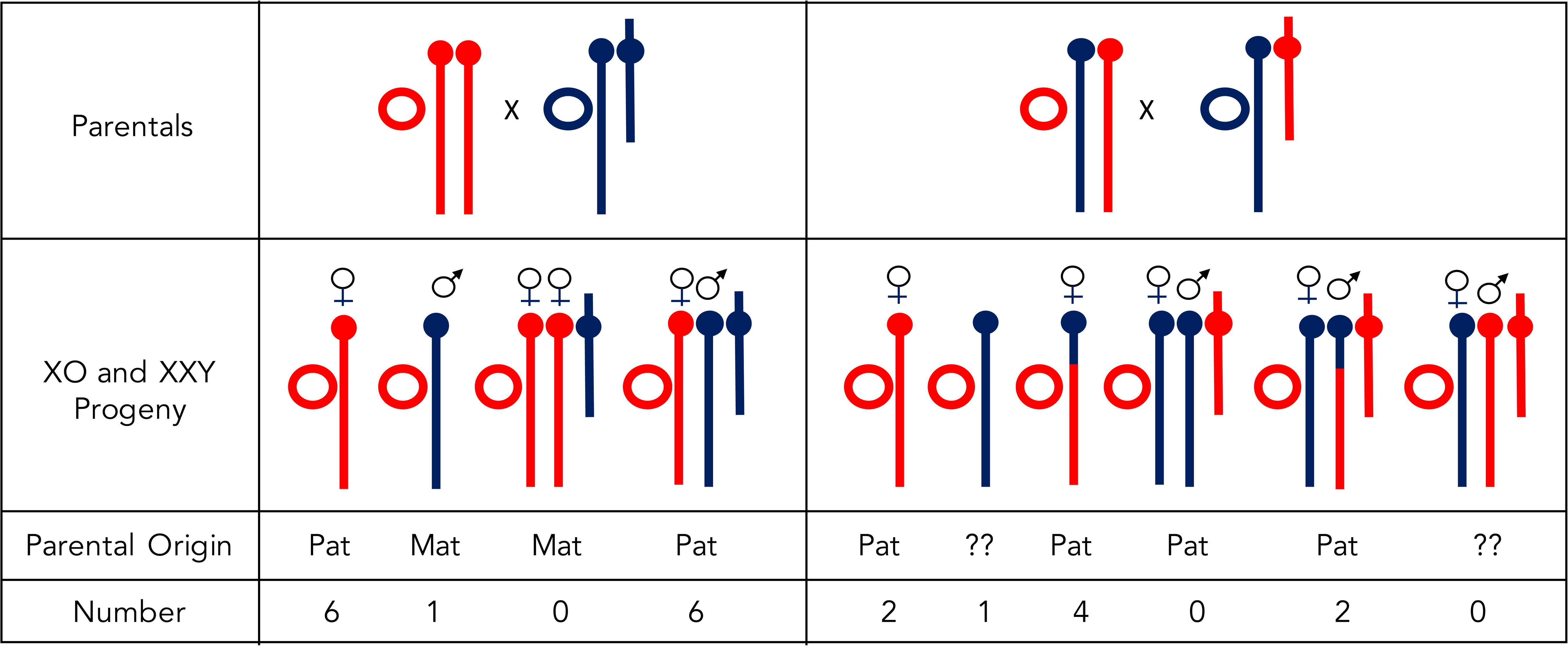
Paternal origin of most sex chromosome aneuploids. Only the sex chromosomes and the mitochondria are shown. The X chromosomes are shown as acrocentric, Y chromosomes as submetacentric and mitochondria as circles. The parents of two types of crosses (outcross or intercross) are shown at the top of the figure with the dam shown on the left and the sire on the right. The potential types of aneuploid progeny in each type of cross are shown with the parental origin below. The figure also shows the inferred parental origin of the aneuploidy and the actual number of those observed in our dataset.

### Detection of sex chromosome mosaicism

There were eight samples (two classified as XX, three as XXY and three as XO) with abnormal chromosome Y intensities (either too low or two high) and with low number of chromosome Y genotype calls (**Figure 1**). Because this pattern suggested mosaicism we performed several additional analyses. As a test case, we selected the tail-derived sample TL9348 (also named Unknown, **Supplementary Tables 1 and 6**) because it was expected to be a F1 hybrid male derived from a C57BL/6J and 129X1/SvJ outcross, has questionable genotype quality and low pd_stat. Based on chromosome intensity this sample was classified as an XXY male with low chromosome Y intensity. Inspection of the genotype calls on chromosome X reveals a significant excess of N calls compared to the autosomes (p<0.00001, **Supplementary Table 6**). Furthermore, the H calls are consistent with the expected contribution of the two parental inbred strains but at only a fraction of expected sites. These results suggest that the mosaicism is due to the loss of both the Y chromosome and one of the two X chromosomes in a fraction of cells. To test this hypothesis, we plotted the intensity of X chromosome markers for three types of controls, C57BL/6J and 129X1/SvJ samples and heterozygous females as well as for the suspected mosaic XXY sample (**Figure 3**). The pattern shown in this figure explain the observed mix of N calls, heterozygous calls and C57BL/6J calls in the XXY sample and confirms its mosaic nature. It further demonstrates that the X chromosome lost is the 129X1/SvJ one. Finally, we can estimate the fraction of cells with XXY and XO constitution using the distance of each maker to their corresponding C57BL/6J and het counterparts. Based on the analysis, we estimate that approximately half of cells are XXY and the other half XO, a result that is also consistent with reduction in the Y chromosome intensity by half. Considered together, these results indicate that the mosaicism occurred early during development, a common observation in embryo mosaicism in humans (Johnson *et al*. 2010; Fragouli *et al*. 2011; McCoy 2017).

**Figure 3.**
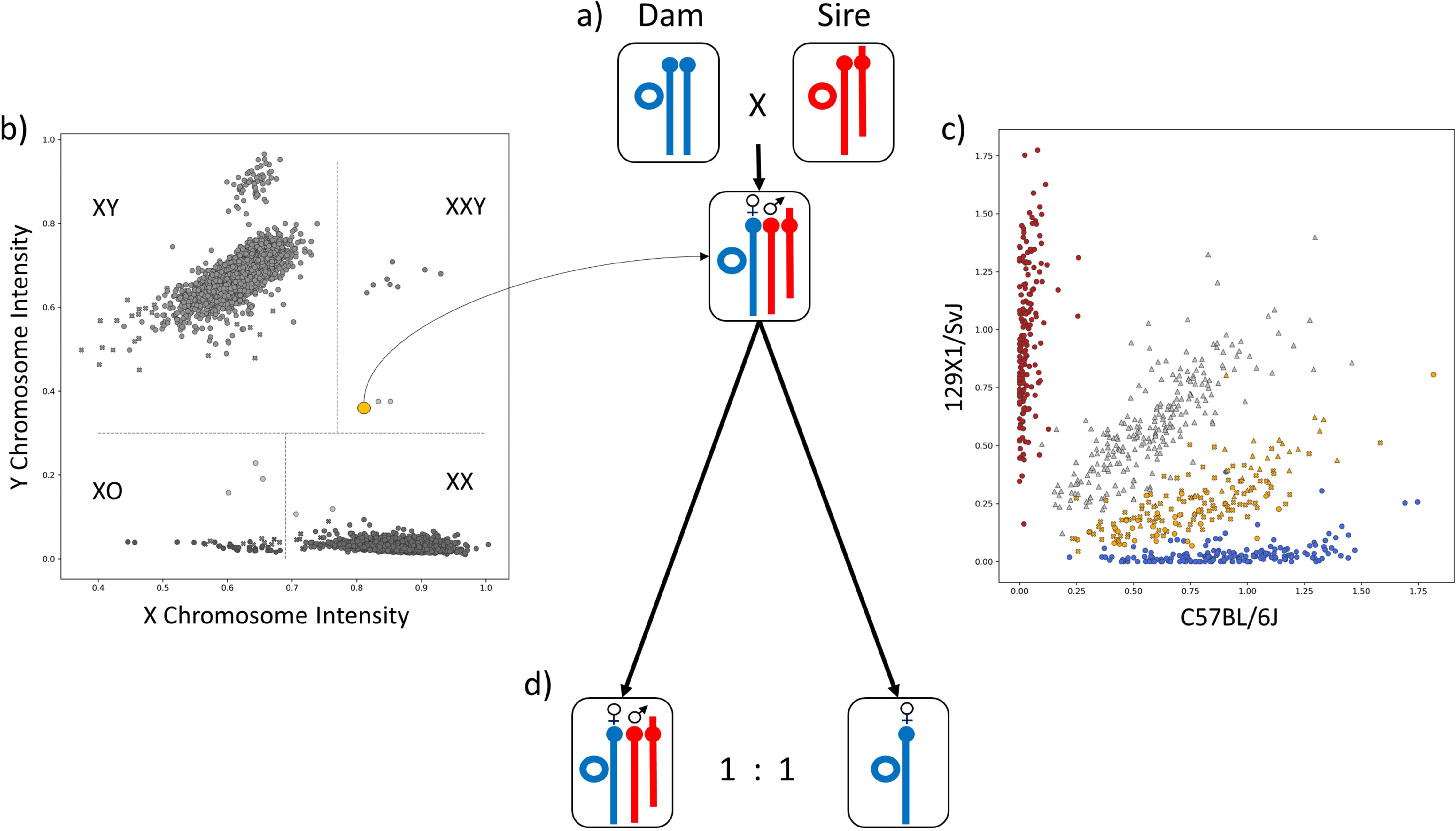
Complex sex chromosome mosaicism in an XXY male. a) shows the chromosomal sex and mitochondria complement of the parents and XXY progeny. b) was used to identify the sex chromosome aneuploidy (two X chromosomes and Y present) and as evidence of mosaicism for presence and absence of Y chromosome (low Y intensity). c) provides evidence of mosaicism for the X chromosome and identifies the paternal origin (129X1/SvJ) of the chromosome lost in some cells. d) The sex chromosome complement of the two types of cells present in this male are shown. Panels b and c were used also to estimate the fraction of each type of cells. (blue points denote C57BL/6J genotype calls, red points 129s1/SvImJ genotype calls. Panels a, c, d).

Among the remaining seven potential mosaics, one was a cell line and thus mosaicism of the sex chromosomes is not unexpected. For the other six samples we performed a similar analysis as the one described above. In all cases the two sets of calls were consistent and thus suggest chromosome Y mosaicism. However only the two samples with 50 or more genotype calls have strong support for such a conclusion. In the Discussion we expand this analysis and provide some guidance for users of the array.

### Strain specific chromosome Y duplications

Among XY males there was a distinct cluster of 64 male samples with higher normalized median Y chromosome intensity (**Figure 1**). These samples include five inbred C3H/HeJ, two F1 hybrid males with a C3H/HeJ chromosome Y (**Figure 4a**) and 52 males derived from a C3H/HeJ by C3H/HeNTac F2 intercross. The plot of the normalized Y chromosome intensity in these males and 81 additional males with Y chromosomes derived from other C3H/He substrains (**Figure 4a**), revealed a clear separation between males carrying a Y chromosome from C3H/HeJ and males carrying C3H/HeNCrl, C3H/HeNHsd, C3H/HeNRj, C3H/HeNTac and C3H/HeOuJ Y chromosomes. Males with the high intensity Y chromosome also include two transgenic strains from The Jackson Laboratory, B6C3-Tg(APPswe,PSEN1dE9)85Dbo/Mmjax and B6;C3-Tg(Prnp-SNCA*A53T)83Vle/J. Both strains were developed and/or maintained in B6C3H background (WEBSITE).

**Figure 4.**
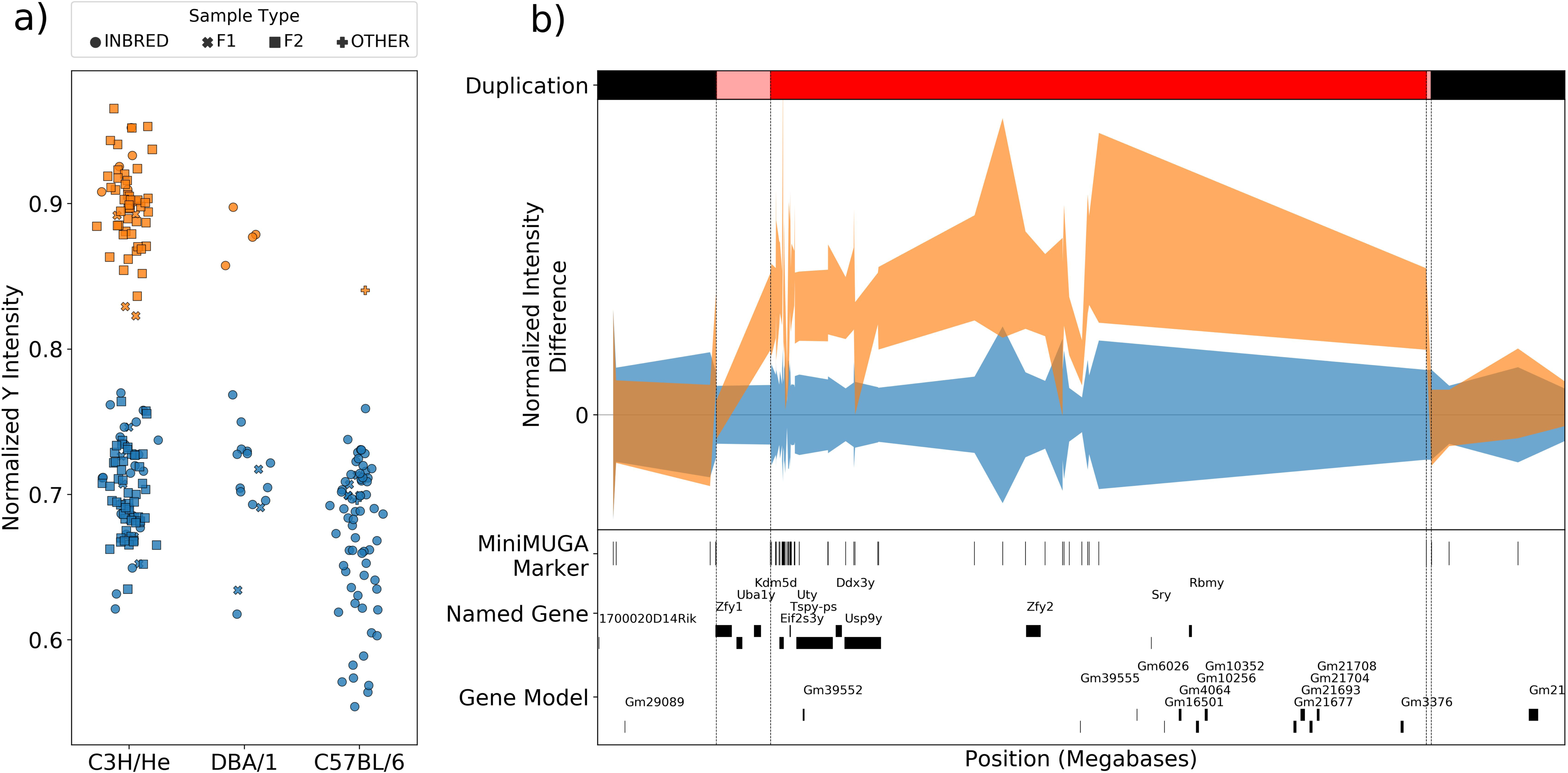
Segmental chromosome Y duplications in laboratory strains. a) Normalized median Y chromosome intensity in selected samples with C3H/He, DBA/1 and C57BL/6 Y chromosomes. Samples with a C3H/HeJ Y chromosome are shown in orange while samples with any other C3H/He Y chromosome are shown in different shades of blue. b) Spatial distribution of normalized intensity at SNPs in the proximal end of the Y chromosome in the same C3H/He samples shown in the a panel. The range of intensities in samples with a C3H/HeJ Y chromosome are shown in orange while samples with any other type of C3H/He Y chromosome are shown in blue. Duplicated region is shown in red and transition regions with uncertain copy number are shown in pink. The bottom of the figure shows the location of the MiniMUGA markers and genes.

To determine the origin of the higher median intensity in males with a C3H/HeJ Y chromosome, we plotted the normalized intensities at MiniMUGA markers located on that chromosome (**Figure 4b**). Inspection of this figure indicates that 54 consecutive markers have distinctly higher intensity and are flanked by markers with intensities that are undistinguishable from males with other C3H/He Y chromosomes. These markers define a 2.9 Mb region located on the short arm of the Y chromosome containing eight known genes *Eif2s3y, Uty, Dxd3y, Usp9y, Zfy2, Sry* and *Rbmy*, and 12 gene models (**Figure 4b**). We conclude that C3H/He substrain differences are due to an intrachromosomal duplication that arose and was fixed in the C3H/HeJ lineage after the isolation of that substrain in 1952 (Akeson *et al*. 2006). There are five additional non-C3H/He samples with high normalized median chromosome Y intensity, four technical replicates from a single DBA/1OlaHsd male and a single *Axl^-/-^* congenic mouse on a C57BL/6 background (**Figure 4a**). Each case represents a different, independent (different haplotype and different boundaries, **Supplementary Figure 5**) and very recent duplication of the Y chromosome. These duplications were segregating within a closed colony. Given that we have identified three independent large segmental duplications of the Y chromosome among 3,018 unique males, we estimate the mutation rate at 1/1000, a relatively high rate. This is consistent with the segmental duplications reported in wild mice (Morgan and Pardo-Manuel de Villena 2017).

### An effective tool for genetic QC in laboratory inbred strains

To determine the performance of MiniMUGA among inbred strains we genotyped 779 samples representing 241 inbred strains including 86 classical inbred strains, 34 wild-derived inbred strains, 49 BXD recombinant inbred lines and 72 CC strains (**Supplementary Table 3**). We created consensus genotypes for each inbred strain using both biological and technical replicates (see Materials and Methods). The use of replicates strengthen the conclusions that can be made from our genetic analyses as they provide a simple but robust method to determine the performance of each SNP in each strain (see Discussion) as well as determining the dates when diagnostic alleles arise and potentially became fixed (see below). We note that for the CC strains, which are incompletely inbred (Srivastava *et al*. 2017), our consensus calls were based on a small number of samples. As such, these consensuses may not completely reflect the individual genotype of any CC animal from a specific strain. Future sampling of a wider range of genotypes from CC mice throughout the history of the CC colony will assist in more accurate consensus genotypes for these strains.

Using the consensus genotypes we determined the number of informative markers for pairwise combinations of all inbred strains. **Figure 5** summarizes the results for 83 classical inbred strains. Over 90% of comparisons have at least 1,280 informative autosomal markers and all but 0.52% of pairwise comparisons have more than 40 informative autosomal markers (2.1 markers per autosome). These statistics are exceptional given the limited number of markers in the array, the inclusion of a large number of diagnostic markers, and a substantial number of construct markers. Although our focus is on classical inbred strains, we extended the analysis to include 37 wild-derived strains. For all 2,924 combinations of classical and wild derived strains, the informativeness is high (mean = 3,224, min = 1,649, max = 3,827). In marked contrast, combinations between wild-derived strains have a much wider range of informative SNPs (from 93 to 3,410) due to a significant fraction of combinations with few to moderate number of informative SNPs. The pairs of strains with the lowest number of informative SNPs include pairs of strain from a taxa other than *Mus musculus* (for example SPRET/EiJ, SMZ and XBS) and pairs of strains that are known to have close phylogenetic relationships (TIRANO/EiJ and ZALENDE/EiJ; and PWD and PWK/PhJ; (Yang *et al*. 2011)). We conclude that MiniMUGA is a significant improvement for genotyping standard lab strains and experimental crosses derived from them.

**Figure 5.**
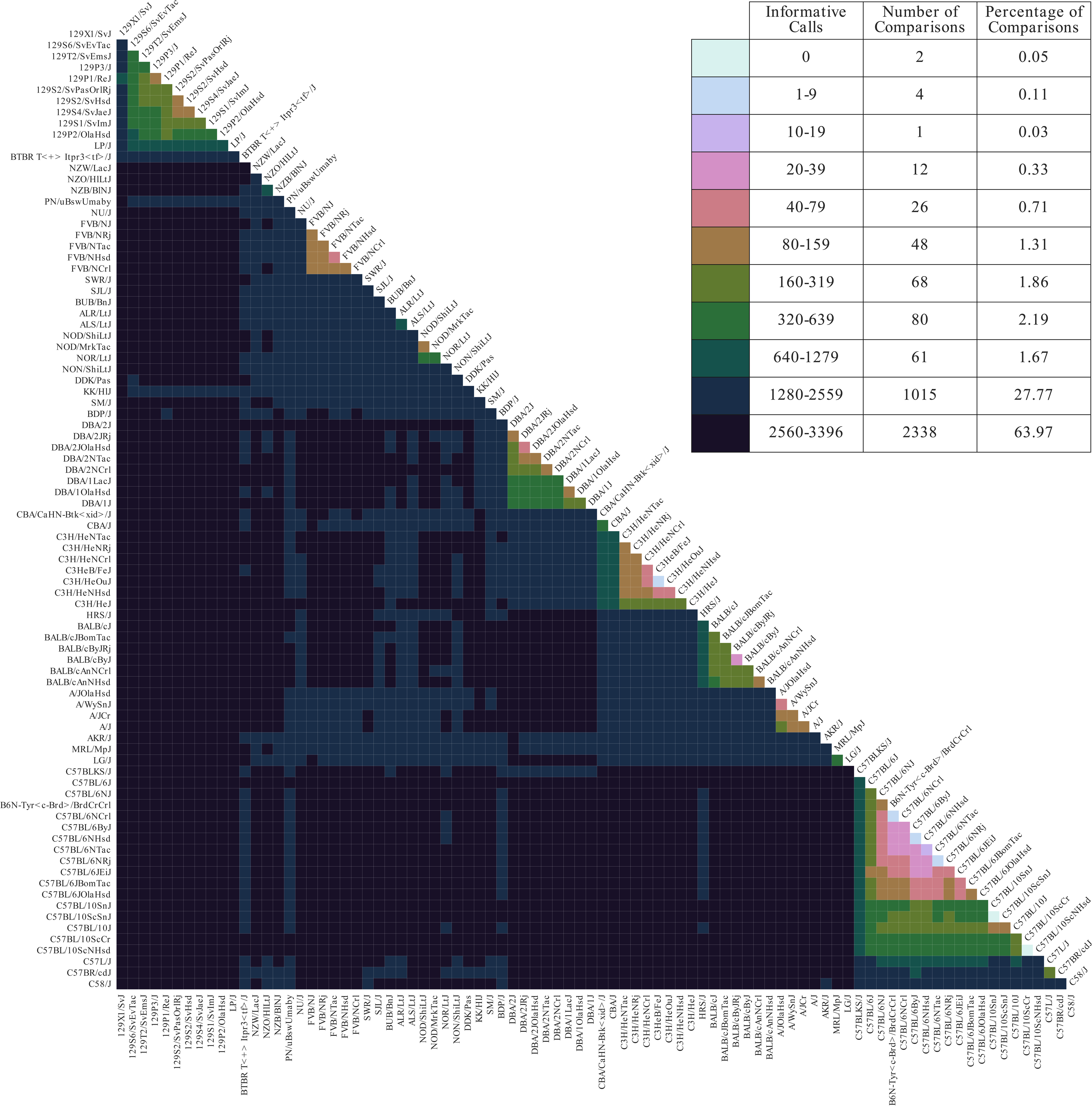
Pairwise number of informative calls in classical inbred strains. Strains are ordered by similarity and colors represent the number of informative SNPs based on the consensus genotypes. Only homozygous base calls, at tier 1 and 2 markers, on the autosomes, X, and PAR are included.

### Mitochondria

MiniMUGA has 88 markers that track the mitochondrial genome, 82 of which segregate in our set of 241 inbred strains. Based on these 82 markers, the inbred strains can be classified into 22 different haplogroups, 19 of which discriminate between *M. musculus* strains (**Figure 6a**). Fifteen haplotypes represent *M. m. domesticus (*groups 1 to 15 in **Figure 6a**), and two haplotypes represent *M. m. musculus* (16 and 17) and two *M. m. castaneus* (18 and 19). Three haplotypes represent different species such as *M. spretus* and *M. macedonicus*.

**Figure 6.**
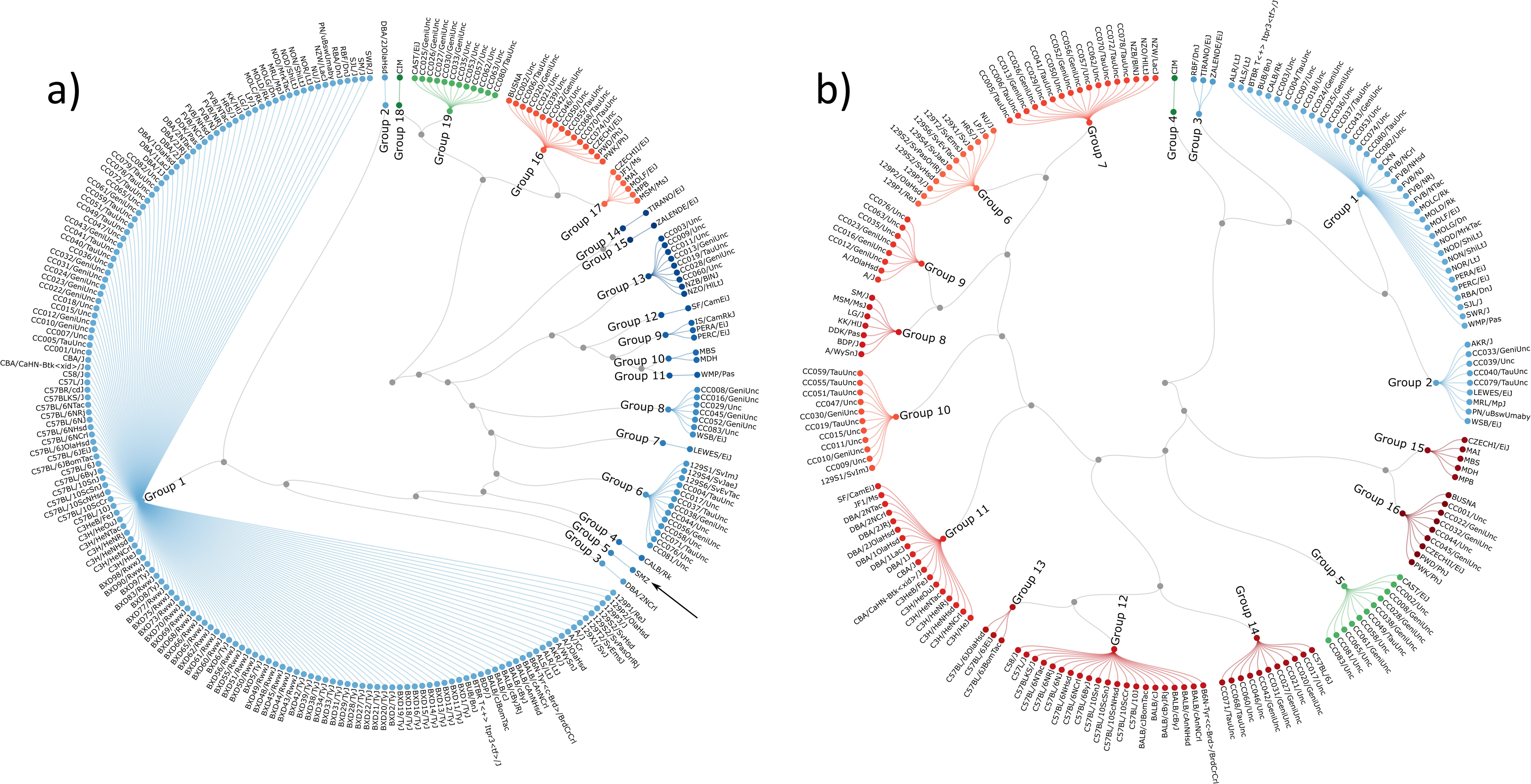
Haplotype diversity of the mitochondria (a) and chromosome Y (b). The trees are built based on the variation present in MiniMUGA and may not represent the real phylogenetic relationships. Colors denote the subspecies-specific origin of the haplotype in question: shades of blue represent *M. m. domesticus* haplotypes; shades of red represent *M. m. musculus* haplotypes; shades of green represent *M. m. castaneus* haplotypes.

In *M. musculus,* nine haplotypes are present in multiple inbred strains while 10 are found in a single inbred strain. The most common haplotype is present in 158 inbred strains (including 49 BXD and 26 CC strains). This haplotype is found in many classical inbred strains including C57BL6/J, BALB/cJ, A/J, C3H/HeJ, DBA/1J, DBA/2J and FVB/NJ. Unique haplotypes represent an interesting mix of wild-derived strains (LEWES/EiJ, CALB/Rk, WMP/Pas, SF/CamEiJ, TIRANO/EiJ, ZALENDE/EiJ, CIM) and DBA/2 substrains (DBA/2JOlaHsd and DBA/2NCrl). CC strains fall into six common haplotypes, one shared by CC three founders A/J, C57BL/6J and NOD/ShiLtJ and five haplotypes present in a single CC founder: PWK/PhJ, 129S1/SvImJ, CAST/EiJ, NZO/HlLtJ and WSB/EiJ. Interestingly, SMZ, a wild-derived inbred strain of *M. spretus* origin, has a mitochondrial haplotype that unambiguously cluster with *M. m. domesticu*s (**Figure 6a**) demonstrating a case of interspecific introgression.

### Chromosome Y

MiniMUGA has 75 markers that track the Y chromosome, 57 of which segregate in our set of 189 inbred strains with at least one male genotyped. Based on these 57 markers, the inbred strains can be classified into 18 different haplogroups, 16 of which are *M. musculus* (**Figure 6b**). Only four haplotypes represent *M. m. domesticus*, two haplotypes represent *M. m. castaneus* and 11 represent *M. m. musculus*. *M. spretus* and *M. macedonicus* are represented by a single haplotype each. In *M. musculus*, all but one haplotype (CIM) are present in multiple inbred strains. No single haplotype dominates in our collection of inbred strains (the most common is present in 38 inbred strains). Interestingly, C57BL/6 substrains fall into three distinct haplotypes. The ancestral haplotype is found in C57BL/6ByJ, C57BL/6NCrl, C57BL/6NHsd, C57BL/6NJ, C57BL/6NRj and B6N-Tyr<C-Brd>/BrdCrCrl. This haplotype is present in other classical inbred strains such as BALB/c, C57BL/10, C57BLKS/J, C57L/J and C58/J. The second haplotype is present in C57BL/6JBomTac, C57BL/6JEiJ and C57BL/6JOlaHsd. Finally, C57BL/6J has its own private haplotype shared with 10 CC strains. Each one of the eight founder strains of the CC (A/J, C57BL/6J, 129S1/SvImJ, NOD/ShiLtJ, NZO/HlLtJ, CAST/EiJ, PWK/PhJ and WSB/EiJ) has its own distinct haplotype.

### Diagnostic SNPs as tool for genetic QC and strain dating

Almost 30% of the SNPs in MiniMUGA are diagnostic for a specific genetic background and were selected from whole genome sequence of 45 classical inbred strains (**Table 2**). We define SNPs as diagnostic when the minor allele is present only in a single classical inbred strain or in a set of closely related substrains. The identification of these SNPs requires WGS from the corresponding strain using the sequence of 12 publicly available strains (Keane *et al*. 2011; Adams *et al*. 2015), 33 substrains that we sequenced and SNP data for the C57BL/10 strain group (**Table 2**). We sequenced these substrains to develop MiniMUGA as well as the desire to expand the number of strains amenable to RCC (MTF, unpub.). Although diagnostic SNPs have low information content (i.e., most samples in a large set of genetically diverse mice will be homozygous for the major allele) they fulfill these two critical missions. First, they increase the specificity of the MiniMUGA array to identify the genetic background present in a sample. In addition, they are essential to extend the power of genetic mapping in RCC beyond the C57BL/6J-C57BL/6NJ paradigm (Kumar *et al*. 2013; Treger *et al*. 2019).

The 3,045 diagnostic SNPs can be divided into two classes based on whether they are diagnostic for a specific substrain (i.e., BALB/cJBomTac or C3H/HeJ) or a strain group (i.e., BALB/c or C3H/He). There are 2,408 SNPs that are diagnostic for one of 45 substrains and 637 SNPs diagnostic for one of 10 strain groups (**Table 2**). A second classification divides diagnostic SNPs into 2,910 fully diagnostic and 129 partially diagnostic SNPs. The difference between these two classes is based on whether the diagnostic allele was fixed or was still segregating in the samples used to determine the consensus genotypes of 46 classical inbred strains.

All diagnostic SNPs started as partially diagnostic SNPs and they highlight the often overlooked fact that mutations arise in all stocks and some of them are fixed despite the best efforts to reduce their frequency and impact. It should be theoretically possible to date when fully and partially diagnostic SNPs arose and whether and when the became fixed in the main stock of an inbred strain. This requires sampling a given substrain at known dates in the past in large enough cohorts to make confident inferences, in other words genotype large cohorts isolated from the main breeding line at known dates.

We have two such populations in our sample set, the BXD and the CC recombinant inbred lines (RIL). In the former we determined whether diagnostic alleles for C57BL/6J and DBA/2J were present in 49 BXD RILs. These RIL were generated in three different epochs: 22 of the genotyped BXD lines belong to epoch I (E1, (Taylor *et al*. 1973)); four belong to epoch II (E2, (Taylor *et al*. 1999)) and 23 belong to epoch III (E3, (Peirce *et al*. 2004)). We determined whether the minor allele at diagnostic SNPs was observed first in epoch I, epoch II, or epoch III. SNPs that were not observed in any of these epochs were grouped under the heading of post E3. **Table 4a** summarizes these findings and further classifies the SNPs based their diagnostic information. We find similar patterns for C57BL/6J and DBA/2J diagnostic SNPs with epoch II contributing the majority of diagnostic SNPs, and epochs III and IV contributing approximately half each of the remaining SNPs. Epoch I SNPs are rare except for the DBA/2 strain group. Finally, and as expected, all partial diagnostic SNPs for C57BL/6J belong to post E3.

**Table 4.**
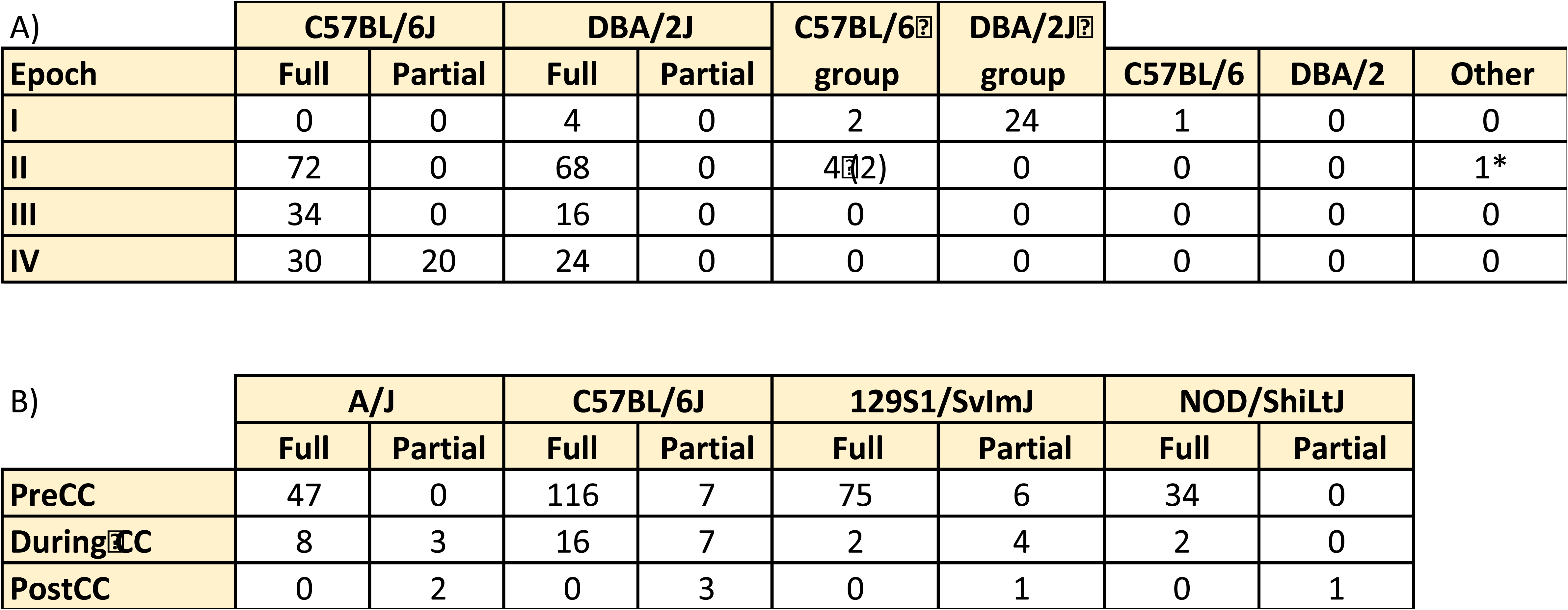

The CC population offers another opportunity to annotate diagnostic SNPs as these RIL were derived from mice from eight inbred strains in 2004 (Collaborative Cross Consortium 2012), including four strains with diagnostics SNPs in MiniMUGA, C57BL/6J, A/J, 129S1/SvImJ and NOD/ShiLtJ. We used 483 CC samples genotyped in the initial array to determine when these 334 diagnostic SNPs arose. We observe three types of patterns depending on the age of the diagnostic allele: 1) the diagnostic allele is fixed in the CC population and thus the diagnostic allele predates the start of the CC project in 2004; 2) the diagnostic allele is absent the CC population and thus the diagnostic allele arose after 2004; and 3) the allele is segregating in the CC with some strains having fixed the diagnostic allele while it is absent in other CC strains. **Table 4b** summarizes these findings.

In addition to determining when diagnostic SNPs arose, it is possible to estimate whether and when the became fixed by examining the allele frequency at consecutive time points and for consistency between populations. This is best exemplified for diagnostic SNPs of the C57BL/6J substrains as we have two time points with substantial sampling, E3 with 23 BXD RIL and the initiation of the CC with 72 CC RIL (note that only one eighth of them will have the C57BL/6J haplotype at any given location and thus the real size of the population used to estimate fixation is closer to 9). There are 75 SNP that were labelled as fixed at E3 because they had 100% allele frequency in both BXD RIL and CC RILs with a C57BL6/J haplotype at the locus. There are also 49 SNPs that were labelled as fixed at the start of the CC because they had 100% allele frequency in CC RILs with a C57BL6/J haplotype at the locus. The remaining 26 diagnostic SNPs were segregating or arose after the start of the CC project. The dates of origin and fixation for diagnostic SNPs are provided in **Supplementary Table 2**.

The birth and fixation of diagnostic alleles can be used to determine the origin and breeding history of a given sample of the appropriate background and thus estimate the expected level of drift (see Discussion).

### Expansion of reduced representation crosses to a large number substrains

We define RCC as crosses between substrains from a single laboratory strain that differ only at mutations that arose after they were isolated and bred independently from the common inbred stock. We tested the ability of MiniMUGA to efficiently cover the genome in 78 different RCC between substrains for which we have consensus genotypes, whole genome sequences and for which live mice are available from commercial vendors (see **Table 2**). We focus our analysis in this group given that WGS of both substrains is required for rapid identification of causative variant(s) (Kumar *et al*. 2013; Treger *et al*. 2019). We used the distance to the nearest informative marker to estimate how well MiniMUGA covers the genome in a given RCC cross. **Figure 7** summarizes these data and demonstrates that for 62 RCCs (82%) all of the genome is covered by a linked marker and in 14 RCCs (18%) between 95% and 99.5% of the genome is covered by a linked marker. Only in two RCCs (3%) there is a significant fraction genome that is not covered by a linked marker. These two crosses are B6N-Tyr<C-Brd>/BrdCrCrl by C57BL/6JOlaHsd and BALB/cByJ by BALB/cByJRj with 8% and 14% of the genome not covered, respectively. An alternative test is the number of RCCs for which 95% of the genome is covered by informative markers at 20cM (56 RCCs or 72%) and 40cM (72 RCCs or 92%) intervals. We conclude that MiniMUGA provides a cost effective tool to extend RCC to substrains from the 129P, 129S, A, BALB/c, C57BL/6, C3H, DBA/1, DBA/2, FVB and NOD strains.

**Figure 7.**
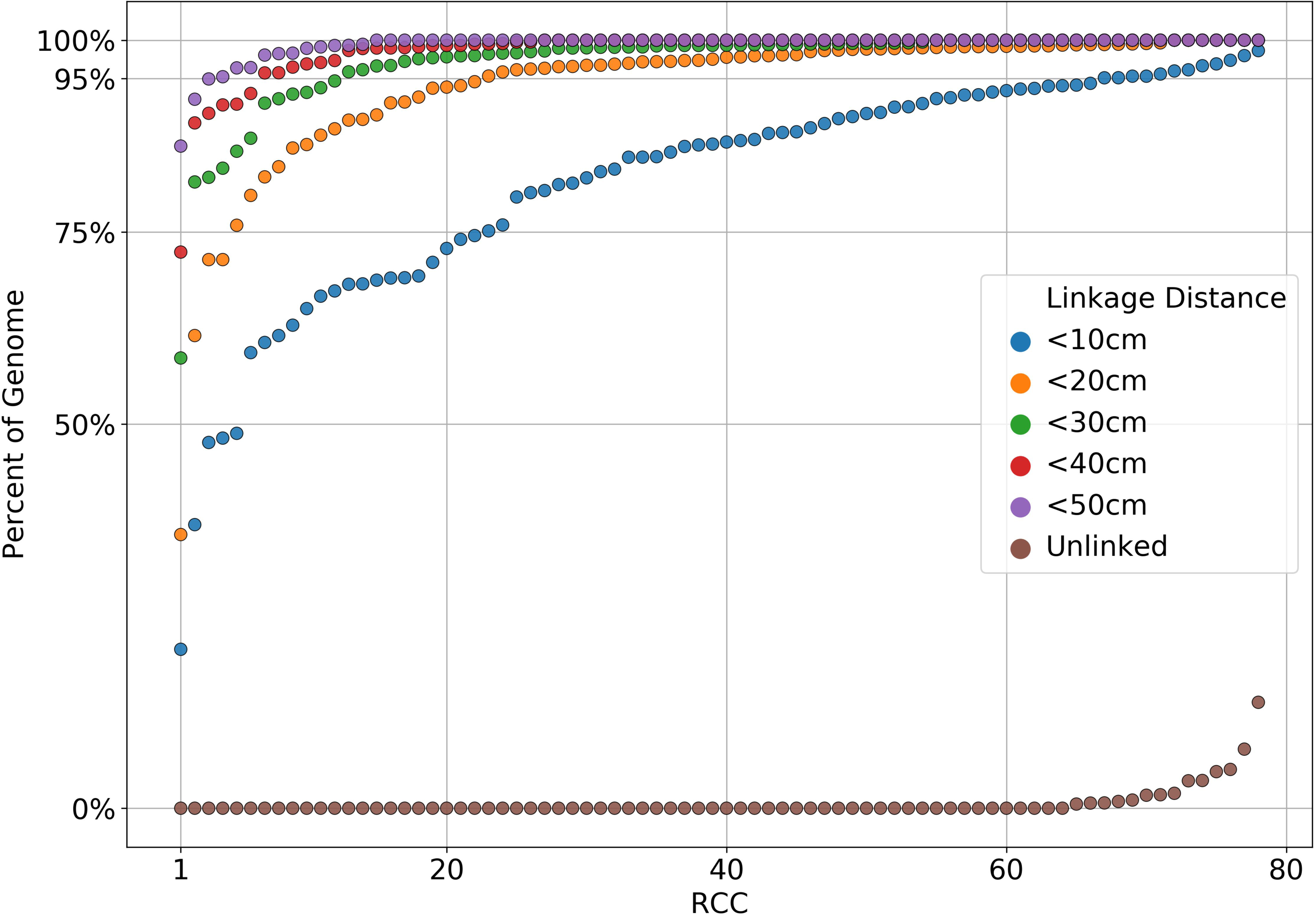
Percent of the genome covered by MiniMUGA in RCCs. The 78 RCCs are shown in ascending order independently for each one of the six analyses. Coverage was based on the linkage distance to the nearest informative marker in a given RCC cross.

### Robust detection of common genetic constructs

Given the broad usage of genetic editing technologies, a key design criterion of MiniMUGA was the ability to detect frequently used genetic constructs. Utilizing our pipeline (low construct probe intensity in classical inbred and F1 samples; variable intensity across the rest of our test population), we positively identified samples containing 17 construct types (**Figure 8**). Importantly, for eight of these constructs, our sample set included positive controls. These positive controls showed robust detection of their relevant constructs. We detected further positive samples from our set in both these eight constructs, as well as nine additional construct classes. All such samples were in sample classes where constructs were plausible (e.g. not wild-derived or CC samples), and there was high concordance for intensities among the probes comprising the detection sets for each of these constructs.

**Figure 8.**
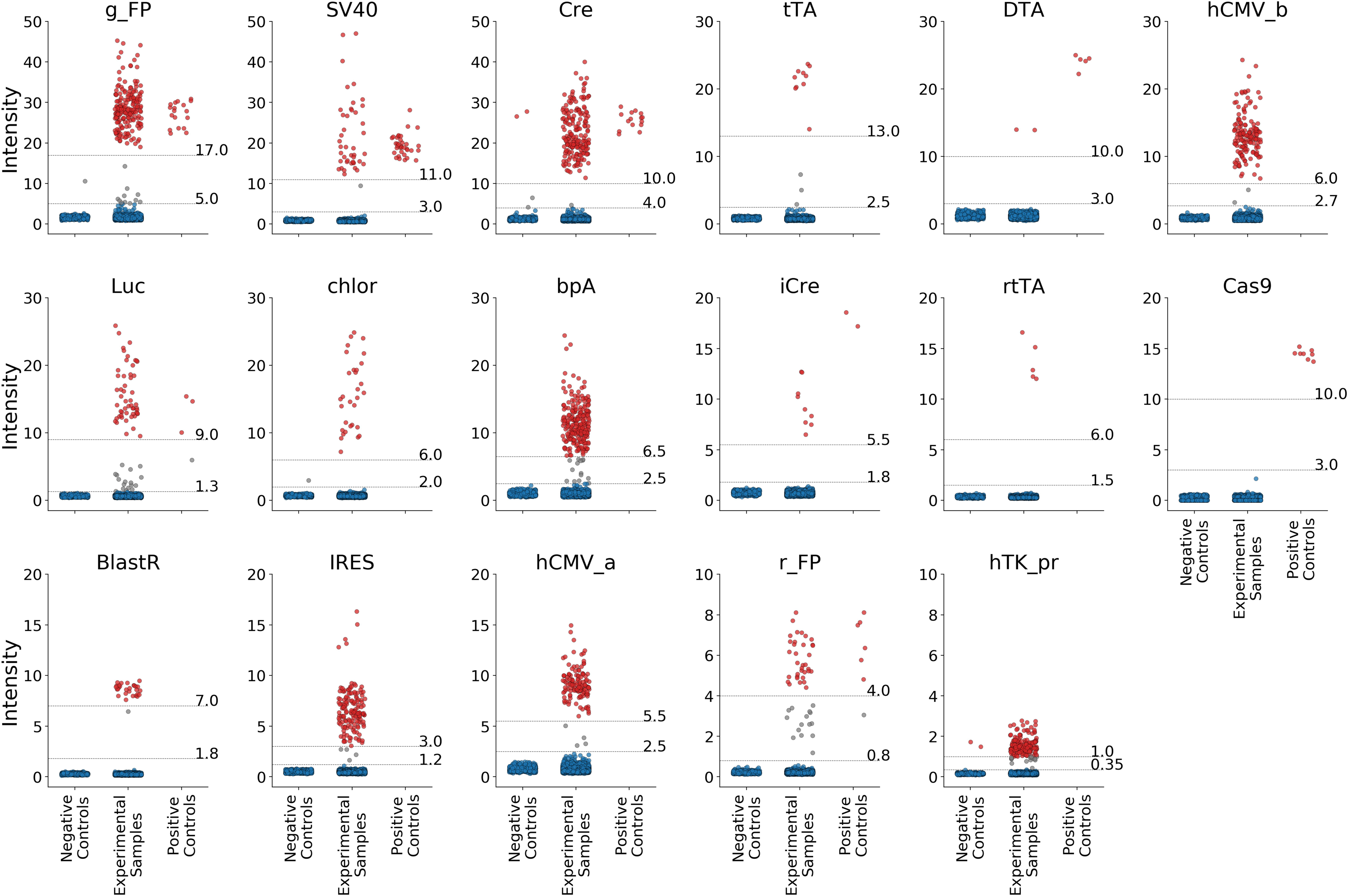
Detection of genetic constructs. For each construct, samples are classified as negative controls (left), experimental (center) and positive controls (right). The dot color denotes whether the sample is determined to be negative (blue), positive (red), or questionable (grey) for the respective construct. For each construct, the grey horizontal lines represent the thresholds for positive and negative results. Note for each construct, the Y-axis scale is different.

For constructs with many probes (**Supplementary Figure 6, Supplementary Table 8**), we noticed that samples we declared as positive could often have significant sample-to-sample variation in their overall intensity (**Figure 8**). As described in the methods and **Supplementary Table 8**, for some construct types our analysis suggested that some probes designed for different constructs were in fact detecting conserved features among multiple construct types (e.g. our ‘g_FP’ designation encompasses probes designed against green-, yellow-, and cyan-fluorescent proteins). As such, it is possible that only a subset of our validated probes are detecting any given sample’s construct. Given our ability to positively identify construct classes with as few as two probes, it is likely that even for constructs which have divergent sequences from our designed sequences, or are targeting a more distantly related construct type, our pipeline will flag samples. An alternative explanation for signal heterogeneity within a construct class is due to within-sample heterogeneity. That is, samples either have variable copies of the construct in question. Such observations might be more common in cell culture samples. Alternatively, construct mosaicism in live animals may manifest as an intermediate signal for given constructs.

As inferred from the above section, across these 17 constructs, we observed that our ability to discriminate between negative and positive samples across these 17 constructs is strongly correlated with the number of independent probes for that construct (**Supplementary Figure 2**, **Figure 8**). As signal intensity is constrained by the dynamic range, our ability to definitively call the presence of low probe number constructs is more uncertain. This uncertainty is especially relevant where a given construct is genetically divergent from the construct sequences used to define a given probe. Users are highly encouraged to consult the probe sequences when they expect a given sample to contain a construct, but do not see support in the array itself. Conversely, for constructs with many independent probes, positive support for a construct is more conclusive, even if a given sample is not expected to contain any constructs.

Finally, we designed probes for 14 constructs, which universally failed in our pipeline. That is, the intensity distributions between known negative (classical inbred strains from commercial vendors and F1 hybrids) and experimental samples were not different. The easiest explanation for these differences is that no samples within our set contained these constructs. Consistent with this explanation is our *a priori* knowledge that no samples in our set could be defined as known positives. In this case, probe-sets may in fact be diagnostic and individual users may identify between sample intensity differences for these constructs. However, as the above sections and methods caution, direct interpretation of single probes or probe-sets are challenging without larger context. Alternatively, though less likely, is that our probe-sets will fail regardless of construct presence. Definitive testing of construct-positive and construct-negative samples for these probe-sets in the future will provide definitive answers to these.

### An easy to interpret report summarizes the genetic QC for every sample

The MiniMUGA Background Analysis Report (**Figure 9**) aims to provide users with essential sample information derived from the genotyping array for every sample genotyped. The report is designed to provide overall sample QC, as well as genetic background information for classical inbred mouse strains, congenic, and transgenic mice. For samples outside of this scope the report may be incomplete or provide misleading conclusions. Details of the thresholds and algorithms for each section of the report are provided in the Materials and Methods section.

**Figure 9.**
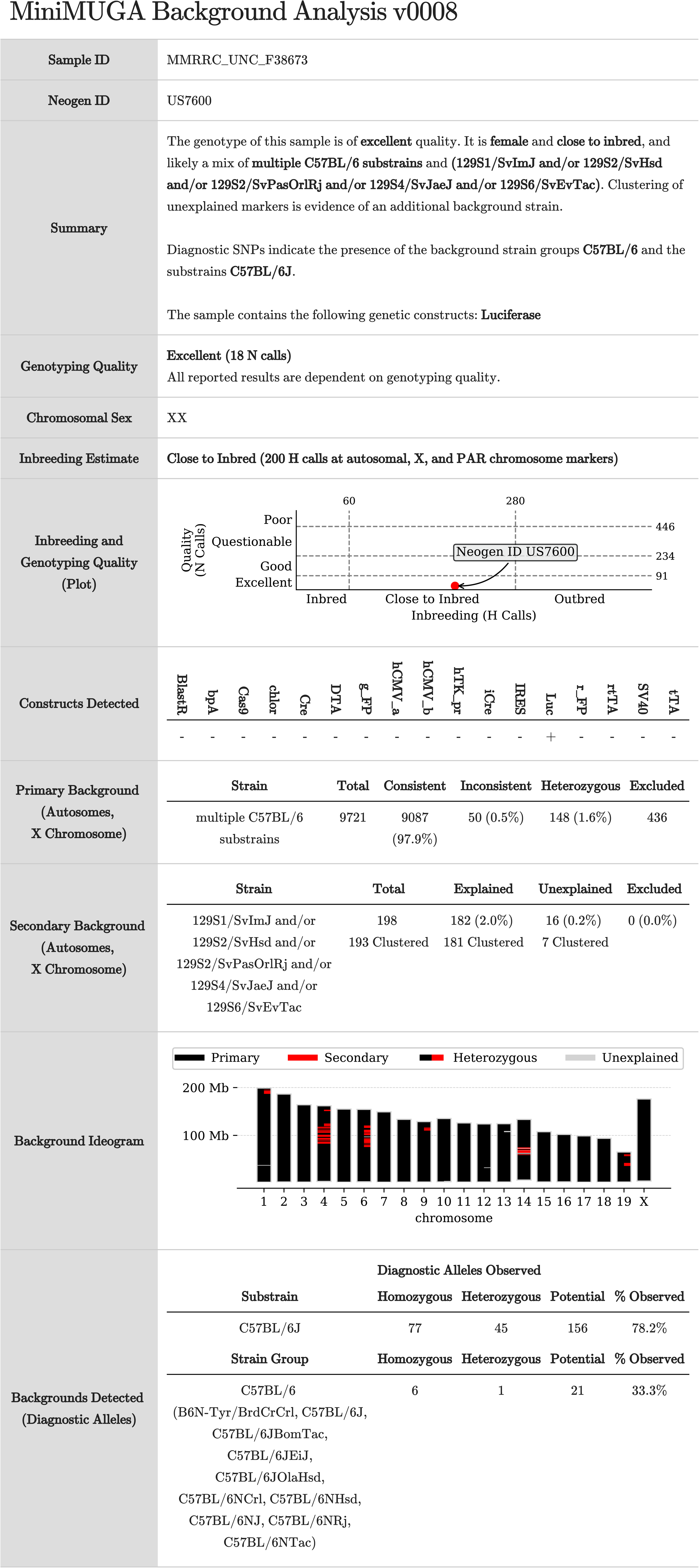
Background Analysis Report for the sample B6.Cg-*Cdkn2a^tm3.1Nesh^ Tyr^c-2J^ Hr^hr^*/Mmnc (named MMRRC_UNC_F38673). The genotype of this sample is of excellent quality. It is a close to inbred female that is a congenic mouse with C57BL/6J as a primary background, and with multiple regions of a 129S background. This sample is positive for a luciferase-family construct and negative for 16 other constructs.

In addition to chromosomal sex and presence of constructs, the report provides a quantitative and qualitative score for genotyping quality. Based on the number of N calls per sample of our sample set we classified samples in one of four categories: samples with Excellent quality (0 to 91 N calls, represents 96.8% of samples); samples with Good quality (between 92 and 234 N calls, 2% of samples), samples with Questionable quality (between 235 and 446 N calls, 0.9% of samples) and samples with Poor quality (more than 447 N calls, 0.3% of samples). Only tier 1 and 2 markers were used in this analysis.

Regarding inbreeding status, the report assigns every sample to one of three categories: Inbred (fewer than 61 H calls), close to inbred (between 61 and 280 H calls) and outbred (more than 280 H calls). These thresholds are based on the number of H calls observed in the autosomes of 172 samples of classical inbred strains and predicted heterozygosity in 3,655 *in silico* F_1_ hybrid mice (**Supplementary Figure 7)**.

For genetic QC, the report provides two complementary analyses. One analysis determines the primary and secondary background of a qualified sample based on the totality of its genotypes (excluding the Y chromosome). The second returns the genetic backgrounds detected in a sample based on the presence of the minor allele at diagnostic SNPs (see section on diagnostic SNPs as tool for genetic QC and strain dating). The initial diagnostic analysis uses the presence of minor alleles in the sample genotypes at identified diagnostic SNPs to identify which (if any) of 46 substrains and/or 10 strain groups are present in the sample.

For the primary background analysis, the sample’s genotype is compared to a set of 120 classical and wild-derived inbred reference strains (**Supplementary Table 3**) to identify the strain that best explains the sample genotypes. If multiple substrains from the same strain group have been detected via diagnostic alleles, or if there is an overrepresentation of a particular diagnostic strain in the unexplained markers, the algorithm generates a composite strain consensus that incorporates all substrains in that strain group and uses it in the primary background analysis. The strain or combination of substrains that best matches the sample is called the primary background for the sample. The report provides the number of homozygous calls that are consistent or inconsistent with the primary background, as well as the number of heterozygous calls in the sample. The primary background is always returned for samples in which the primary background explains at least 99.8% of the sample genotype calls.

Once the primary background is determined, the algorithm tests whether at least 75% of the markers inconsistent with the primary strain background or heterozygous (aka unexplained) are spatially clustered. If they are not (<75% of markers spatially clustered) the algorithm will not try to identify a secondary background. If at least 75% of the unexplained markers are clustered, all strain(s) from the reference set that best explain at least half of the unexplained calls are identified as secondary background(s). If the combination of primary and secondary backgrounds explains at least 99.8% of the calls, the primary and secondary backgrounds are reported. If it explains <99.8% then no genetic background is returned.

For samples where a primary and secondary background is reported, the algorithm determines whether the remaining unexplained markers are spatially clustered. If they are, the summary states that clustering of unexplained markers may indicate the presence of an additional genetic background. The limitations of this greedy approach to identification of the primary and secondary backgrounds are further explained in the Discussion section.

Note that this report is generated programmatically using the available reference inbred strains (**Supplementary Table 3**). If the reported results are inconsistent with expectations, users need to consider further analyses before reaching a final conclusion. All estimates and claims in the report are heavily dependent on the quality of the sample and genotyping results. Less than excellent genotyping quality will likely increase the likelihood of an incorrect conclusion. Genotyping noise can lead to incorrect reporting and may be particularly misleading in samples from standard commercial inbred strains. Fully inbred strains routinely have a small percentage of spurious H calls. These do not represent true heterozygosity (see consensus of inbred strains).

### Cell lines

Cell lines can be subject to the same genetic QC as mice. We have previously reported that the number of N calls is higher for cell lines that mixed tissues in other arrays (Didion *et al*. 2014). There is some evidence of this in our dataset but it is inconclusive. We have already shown the ability to detect sex chromosome aneuploidy in cell lines (**Figure 1**). Diagnostic SNPs can be use to date cell lines in similar fashion with the added simplicity that cell lines are less susceptible to change. Finally, cell line can be run the same Background Analysis Report pipeline discussed in the previous section, some examples are provided in **Supplementary Figure 8**. The importance of genetic QC in cell lines will grow in future given the increased emphasis on cell based research.

## DISCUSSION

### MiniMUGA as a tool for QC

Among the many new capabilities of the MiniMUGA array compared with its predecessors is the Background Analysis Report provided with each genotyped sample. Although expert users can, and undoubtedly will, refine existing and develop new analyses pipelines; all users benefit from a common baseline developed after the analyses of many thousands of samples. The size, annotation, and variety of our sample set provided a firm foundation for the results and conclusions presented here.

We urge users to pay particular attention to genotype quality, reported heterozygosity and unexpected conclusions (i.e., sex, backgrounds and constructs detected). Genotype quality depends on the sample quality, quantity and purity and on the actual genotyping process. Poor sample quality can also be the byproduct of off target variants in the probes used for genotyping and thus wild mouse samples and mice from related taxa are expected to have lower apparent quality. Samples with poor quality will not be run through the report. Samples with questionable quality may lead to incorrect conclusions. For samples of any quality the total number of N calls should be carefully considered if unexpected results are reported. It is also important to consider the pd_stat when evaluating the chromosomal sex determination.

Reported heterozygosity is sensitive to genotyping quality. A lower quality sample will typically include more spurious heterozygous calls than an excellent quality sample of the same strain. This leads to an incorrect estimate of the level of inbreeding in a given sample, and can be particularly misleading in a fully inbred mouse of pure background. The thresholds used to classify samples as inbred, close to inbred and outbred are somewhat arbitrary and reflect the biases in SNP selection (overrepresentation of diagnostic SNPs for selected substrains) and the highly variable range of diversity observed in F1 mice. We used the observed number of H calls in known inbred samples and the predicted number of H calls among a large and varied set of potential F1 hybrids to set our thresholds, but users should define the level of heterozygosity in a specific experiment (**Supplementary Figure 7**). For example, mice generated in RCCs between related substrains may have a very small number of H calls and thus will be misclassified as more inbred that they really are. The report combines sample quality and heterozygosity in a single figure for quick visual inspection. Note that the x and y axes are compressed in the high value range to ensure that all samples, even those with very poor quality and/or high heterozygosity, are shown. The precise location of a sample in the plot should help customers contextualize their sample’s quality and inbreeding when evaluating their results.

For users genotyping large number of samples in a given batch (for example, several 96 well plates) we found it useful to include a plate-specific control at an unambiguous location (we use the B3 well). Ideally, these controls have known genotypes, excellent quality and are easy to distinguish from all other samples in the batch. Plating errors or unaccounted transpositions occurring during the genotyping process are rare but problematic. Adding one sample per plate is a reasonable cost to quickly identify these issues before they metastasize.

We anticipate that most users will use the Background Analysis Report to determine the genetic background(s) present in a sample as well as their respective contributions. The identification of the correct primary and secondary background is completely dependent on the pre-existing set of reference strains (**Supplementary Table 3**). If a genotyped sample is derived from a strain that is not part of this reference set, the reported results may be misleading or completely incorrect. Users should consult the list of reference backgrounds. We expect the number of reference backgrounds to increase over time, reducing the frequency and impact of this problem. However, the current background detection pipeline is not appropriate for recombinant inbred lines (RIL) such as the BXD and CC populations. By their very nature RIL have mosaic genomes derived from two or more inbred strains (included in our panel) and thus the background analysis will detect more than two inbred backgrounds (for CC strains) or declare one of the parental strains as a secondary background in some specific cases. Users interested in confirming or determining the identity of RIL can use our consensus genotypes to do so.

An important caveat of the current primary and secondary background analysis is that the approach is greedy, and all variants except those with H and N calls in the consensus are considered. Because only a fraction of the SNPs are informative between a given pair of strains (typically less than half, see **Figure 5** and **Supplementary Figure 7**), the algorithm always overestimates the contribution of the primary background and underestimates the contribution of the secondary background (**Figure 9**). As a general rule in congenic strains, the contribution of the strain identified as the secondary background should be multiplied by at least 3. If the exact contribution of either background is critical for the research question, the user should reanalyze the data using only SNPs that discriminate between the two backgrounds.

A second caveat is that the current pipeline does not include the mitochondria and Y chromosomes’ genome. This shortcoming will be addressed in a future update of the Background Analysis Report.

A final caveat is that in most cases if more than two inbred strains are needed to explain the genotypes of a sample, the report does not identify any of them. In our experience when three of more backgrounds are present a greedy search is not effective and often leads to incorrect results. If the user has prior knowledge of at least some of the backgrounds involved, an iterative hierarchical search will typically yield the correct solution, but care needs to be taken at each step.

Genetic constructs have been a staple of genome editing technologies since the 1980s. In addition to desired genetic modifications, constructs will often include a variety of other necessary features (e.g. selection markers; constitutive promoters). The array can be used to validate the presence of constructs expected to be present and/or to identify unexpected constructs.

Our construct probe design was focused on targeting conserved features of various genetic engineering and/or in vitro constructs commonly used in mammalian genetics. We can split these conserved probe-sets into two main classes: those for which we were able to detect positive samples in this large cohort, and those for which we were not able to detect any consistently positive signal/sample. Many of the probe-sets that are reported jointly as a single construct type because the signals were highly correlated (e.g. the cyan, green and yellow fluorescent protein probe-sets). Interested users can use the individual probe intensities to refine the analysis.

Similarly, in the dataset used to define the performance of the array, we were unable to identify samples positive for several individual probes and even some entire probe-sets (**Supplemental Figure 6)**. In some cases we excluded probes due to the fact that they work for different subsets of samples than the included probes (see hTK_pr in **Supplementary Figure 6**). In other cases, the excluded probes failed for an unknown reason and likely cannot be rescued (iCRE in **Supplementary Figure 6)**. Finally, addition of known positive controls may allow the rescue of one or more of the 13 constructs targeted (e.g. ampicillin resistance **Supplementary Figure 6**).

### MiniMUGA as a tool for discovery

MiniMUGA was designed to support the research mission of geneticists, but the range of applications will depend on the ingenuity of its users. In the results sections we explored three areas in which MiniMUGA has high potential to complement existing resources and tools.

The first of these areas is sex chromosome biology. MiniMUGA is able to robustly determine four sex chromosome configurations (**Figure 1**) and thus facilitates estimation of the incidence and prevalence of sex chromosome aneuploidy in the mouse. The variation of aneuploidy rates depending on the sire background provides a promising avenue to study the genetics of sex chromosome missegregation. In addition, identification of aneuploid mice can become routine in experimental cohorts and crosses. This is also important in colony management, as XO and XXY mice are likely to be infertile or have reduced fertility (Heard and Turner 2011).

This type of analysis can also identify sex chromosome mosaicism (Johnson *et al*. 2010; Fragouli *et al*. 2011; McCoy 2017) and large structural variants involving the sex chromosomes. In the results section we have shown that mosaics are outliers from the four defined clusters observed in the intensity based chromosome sex determination plot (**Figure 1**). Specifically, they have abnormal Y chromosome intensities. These mosaics may also have an abnormally high ratio of N calls in the X chromosome compared to the autosomes and chromosome X marker intensity distributions biased towards one parent (**Figure 3**). The last analyses are only possible in the presence of heterozygosity on the X chromosome.

In addition, MiniMUGA revealed a 6Mb *de novo* duplication of the distal chromosome X (**Supplementary Figure 9**) in an F2 male. The size of this duplication is not large enough to affect chromosomal sex determination and its discovery was due to the presence of 10 heterozygous calls clustered on distal X. These heterozygous calls occur at informative markers between the two CC strains involved in the F2 cross and are embedded in a region of 26 consecutive markers with higher than expected intensity (**Supplementary Figure 9**). Interestingly, the parental CC strains (CC029/Unc and CC030/GeniUnc) are the same for which a 10X increase in sex chromosome aneuploidy is observed. We concluded that this F2 male had a sharply defined duplication of the distal X chromosome. These vignettes provide a potential blueprint that can be extended to other chromosomes and structural variants. It also highlights the importance of having a large set of well-defined genotyped controls, against which to compare a given sample.

A second area of potential research is the expansion of the RCC paradigm beyond the narrow confines of C57BL/6N (Kumar *et al*. 2013; Babbs *et al*. 2019; Treger *et al*. 2019). A successful RCC requires complete knowledge of the sequence variants shared and private to the set of substrains that will be used in the mapping experiments. These private variants are obviously needed to infer causation but also in the initial step of genetic mapping. We acknowledge that the development of MiniMUGA was made possible by the efforts of the community to sequence an increasing number of inbred strains. The expansion of RCCs to 129S, A, BALB/c, C57BL/6, C3H, DBA/1, DBA/2, FVB and NOD substrains should increase the total number of accessible private mutations by at least one order of magnitude; and therefore, we should expect a similar increase in the number of mappable causative genetic variants for biomedical traits. We note that even as substrains continue to accumulate private variants in an unpredictable manner, MiniMUGA will retain its value for genetic mapping but additional WGS will be required.

Finally, the private variants that underlie the RCC concept are the diagnostic variants used in background determination and sample dating. Diagnostic SNPs have little information content but high specificity. The presence of diagnostic alleles in a sample is strong evidence that that specific substrain (or a closely related substrain absent from our set) contributed to the genetic background of that sample. However, because only a small fraction of diagnostic SNPs have been observed in all three genotypes across multiple samples, their performance is not well established, in particular for heterozygous calls. To avoid errors, we required diagnostic alleles at three different SNPs in a given sample before a genetic background is declared in the Background Analysis Report. All diagnostic SNPs began their history as partially diagnostic (segregating in an inbred strain or substrain population).

To test whether it is possible to use the annotated diagnostic SNPs to determine the age and breeding history of a given sample or stock we selected 485 samples that where over inbred, had over 99% identity to C57BL/6J and had no diagnostic alleles for any other substrain. The analysis is based in the pattern of ancestral diagnostic SNPs that are classified as fixed in epoch III (E3) or prior to the CC based. **Figure 10** shows three examples with different patterns. Panel A shows a KO mouse from line created prior to epoch III (E3) and bred independently from the C57BL/6J stock since at least 2004. The former conclusion is based the fact that we detect the ancestral allele at 21 SNPs that were fixed prior to epoch III. The later is based in the observation of ancestral alleles at 36 SNPs that we believe to be fixed by 2004 (21 and 15 from E3 and CC, respectively) and that these markers are distributed across 14 chromosomes. Panel B shows a transgenic mouse from a line created prior to the initiation of the CC (2004) and bred independently from the C57BL/6J stock since them. Both conclusions are based the fact that there are zero ancestral alleles at any of 75 diagnostic SNPs fixed by epoch III, the detection of the ancestral allele at 18 SNPs that were fixed prior to the CC and that these markers are distributed across 13 chromosomes. Finally, panel C shows a wild type C57BL/6J mouse derived from the JAX colony after 2004. The conclusion is based in the lack of ancestral alleles at any of 124 fixed diagnostic SNPs and the presence of a derived allele at three SNPs that arose after the CC. Notably our conclusions were consistent with the expectations from the owners of these samples. However, these are fairly simple examples but more complex and more interesting patterns are plentiful in our dataset. For example, four samples from a congenic inbred stock show evidence of both an old stock and new refreshing of the genome in recent years (**Supplementary Figure 11**). Specifically, the presence of ancestral alleles at many diagnostic SNPs fixed prior to epoch III and the start of the CC speaks of mouse line generated and bred independently for many years. On the other hand, heterozygosity at some of these markers as well as the presence of post CC diagnostic alleles indicates that this line we refreshed by backcrossing to C57BL/6J in recent years. Both conclusions are correct as this stock was imported by Mark Heise at UNC in 2014 and backcrossed once or a few times to JAX mice before being maintained by brother sister mating. In addition to improving the genetic QC, we believe that this type of analysis may provide researchers with critical information to guide both experimental design and data analysis. Most important is the ability to estimate the amount of drift that has taken and thus the amount of genetic variation present in that line but absent in the main stock. We expect that use of MiniMUGA and the continued and rapid annotation of diagnostic SNPs not only for C57BL/6J but for all inbred substrains offers an opportunity to significantly improve the rigor and reproducibility of mouse research.

**Figure 10.**
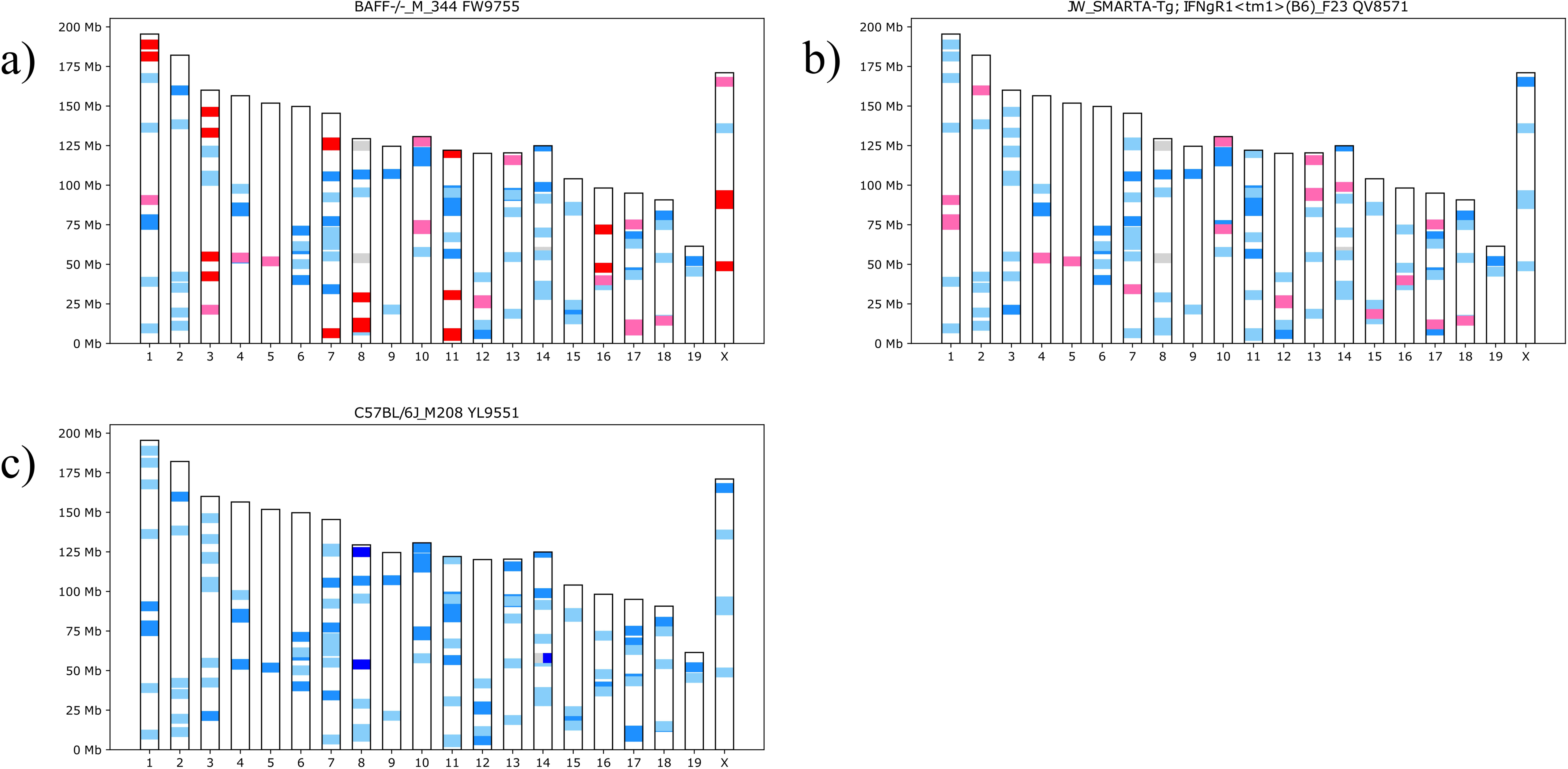
Age and breeding history of mouse samples with C57BL/6J background. a) Inbred Baff^-/-^ male in C57BL/6J background. b) Inbred transgenic and IFNgR1 female in C57BL/6J background. c) Inbred C57BL/6J male. Red bars denote the ancestral allele for diagnostic SNPs fixed at E3. Pink bars denote ancestral alleles for diagnostic SNPs fixed at the start of the CC. Light blue bars denote diagnostic alleles at diagnostic SNPs fixed at E3. Lighter blue bars denote diagnostic alleles at diagnostic SNPs fixed at start of CC. Grey bars denote ancestral alleles at post-CC diagnostic SNPs. Dark blue bars denote diagnostic alleles at post-CC diagnostic SNPs. Split bars denote heterozygous SNPs in a sample.

## Acknowledgments

This work was supported in part by U24HG010100 (to LM and FPMV); U42OD010924 (to TM); U19AI100625 (to RSB, MTH, FPMV and MTF) and P01AI132130 (to CS, MTF and FPMV); R01GM121806 (to JMC), P50DA039841 (to LT), R01MH100241 (to LT, WV, and FPMV), 5R01HL128119 and 5R01DK058702 (to TK), and U42 OD010921 (to CL and LR). The array was used for authentication of key biological materials in the following grants: R01ES029925 and P42ES031007 (to FPMV). The Systems Genetics Core Facility and Mutant Mouse Resource and Research Center at the University of North Carolina provided in kind resources. MiniMUGA was developed under a service contract to FPMV and LM from Neogen Inc., Lincoln, NE. None of the authors have a financial relationship with Neogen Inc. apart from the service contract listed above. The authors have no other conflict of interest to declare. We would like to acknowledge Mohanish Deshmukh, Bev Koller, Helen Lazear, Lawrence E. Ostrowski, and Patrick Sullivan for kindly providing some of the samples.

## SUPPLEMENTARY MATERIAL LEGENDS

**Supplementary Table 1.** Samples included in this study. The table provides the following information:

<u>Serial ID</u>.

<u>Sample name</u>: name provided by the investigator.

<u>Type</u>: inbred, F1, cell line, cross or unclassified.

<u>Content Type</u>: initial or Final.

<u>Consensus strain</u>: if a sample was used to build the consensus genotypes of one 241 inbred strain, that strain name is listed, if that sample was not used then zero.

<u>Chromosomal sex marker selection</u>: TRUE for samples used in selecting sex informative markers. FALSE for samples not used.

<u>Chromosomal sex</u>: XX, XY, X0, XXY or XX*. The latter group are XX samples misclassified as XO.

<u>Replicate</u>: TRUE for technical replicates genotyped more than once. FALSE for samples genotyped only once.

<u>Replicate name</u>: An unambiguous name for that group of replicate samples.

<u>X chromosome inten</u>sity: median normalized intensity of chromosome X sex-informative markers.

<u>Y chromosome intensity</u>: median normalized intensity of chromosome Y sex-informative markers.

<u>Median autosomal intensity</u>: median intensity of autosomal markers (i.e., normalization factor)

<u>H calls</u>: Number of heterozygous calls for tier 1 and 2 markers (see below) in the autosomes and chromosome X.

<u>H call on chromosome X</u>: Number of heterozygous calls for tier 1 and 2 markers (see below) on chromosome X.

<u>Autosomal N calls</u>: Number of no calls for tier 1 and 2 markers (see below) in the autosomes.

<u>N calls on chromosome X</u>: Number of no calls for tier 1 and 2 markers (see below) in the X chromosome.

<u>ks_stat</u>: Kolmogorov-Smirnov goodness of fit test statistic of the sample’s autosomal intensities against the autosomal intensity distribution of 200 random samples

<u>pd_stat</u>: Pearson’s chi-squared test statistic of the sample’s autosomal intensities against the autosomal intensity distribution of 200 random samples.

<u>BlastR</u>: Sum of the autosome-normalized xraw intensity at 6 markers used to declare the presence or absence of the construct Blasticidin resistance.

<u>Cas9</u>: Sum of the autosome-normalized xraw intensity at 7 markers used to declare the presence or absence of the construct Cas9

<u>Cre</u>: Sum of the autosome-normalized xraw intensity at 15 markers used to declare the presence or absence of the construct Cre recombinase

<u>DT</u>: Sum of the autosome-normalized xraw intensity at 11 markers used to declare the presence or absence of the construct Diptheria toxin

<u>IRES</u>: Sum of the autosome-normalized xraw intensity at 6 markers used to declare the presence or absence of the construct Internal Ribosome Entry Site

<u>Luc</u>: Sum of the autosome-normalized xraw intensity at 10 markers used to declare the presence or absence of the construct Luciferase

<u>SV40</u>: Sum of the autosome-normalized xraw intensity at 18 markers used to declare the presence or absence of the construct SV40 large T antigen

<u>bpA</u>: Sum of the autosome-normalized xraw intensity at 8 markers used to declare the presence or absence of the construct Bovine growth hormone poly A signal sequence

<u>chlor</u>: Sum of the autosome-normalized xraw intensity at 9 markers used to declare the presence or absence of the construct Chloramphenicol acetyltransferase

<u>g FP</u>: Sum of autosome-normalized xraw intensity at 19 markers used to declare the presence or absence of the construct “Greenish” Fluorescent Protein (EGFP, EYFP, ECFP)

<u>hCMV a</u>: Sum of the autosome-normalized xraw intensity at 6 markers used to declare the presence or absence of the construct hCMV enhancer version a.

<u>hCMV b</u>: Sum of the autosome-normalized xraw intensity at 11 markers used to declare the presence or absence of the construct hCMV enhancer version b

<u>hTK pr</u>: Sum of the autosome-normalized xraw intensity at 2 markers used to declare the presence or absence of the construct Herpesvirus TK promoter

<u>iCre</u>: Sum of the autosome-normalized xraw intensity at 8 markers used to declare the presence or absence of the construct iCre recombinase

<u>r FP</u>: Sum of the autosome-normalized xraw intensity at 5 markers used to declare the presence or absence of the construct “Reddish” fluorescent protein (tdTomato, mCherry)

<u>rtTA</u>: Sum of the autosome-normalized xraw intensity at 8 markers used to declare the presence or absence of the construct Reverse improved tetracycline-controlled transactivator

<u>tTA</u>: Sum of the autosome-normalized xraw intensity at 14 markers used to declare the presence or absence of the construct Tetracycline repressor protein

**Supplementary Table 2.** Marker annotation. The table contains the following information:

1) Marker name.

2) Chromosome. The following types are allowed: 1-19, for the autosomes; X and Y for the sex chromosomes; PAR, for markers on the pseudoautosomal region; MT, for the mitochondria and 0, for genetic constructs.

3) Position in bases in build 38.

4) Strand. +, indicating the probe sequence is found in the 5’ to 3’ order (on the forward strand) in the reference genome immediately preceding the variant. -, indicating that the reverse complement of the probe sequence is found in the 5’ to 3’ order (on the forward strand) in the reference genome, immediately following the variant and NA, when not available.

5-6) <u>Sequences A and B</u>. Sequence A for one bead probes is the sequence of the marker probe without the SNP and for two bead probes, the sequence of the marker probe including the SNP. Sequence B: for one bead probes, not applicable; for two bead probes, the alternative sequence of the marker probe including the SNP.

7-8) <u>Reference Allele and Alternate allele</u>. Columns denoting the genotype call for the reference and alternative alleles

9) Tier. For biallelic SNP markers, tier was assigned based on observed genotype call types (homozygous reference, homozygous alternate, or heterozygous) at each marker across a set of 3,878 samples used for array QC and validation. Tier 1 markers were those for which we observe all three call types. Tier 2 markers were those for which we observe two of the three call types. Tier 3 markers were those for which we observe only one call type. Tier 4 markers were those markers for which we observe no calls (N) in every sample. For construct markers, tier is assigned based on the capability of a marker to detect a given construct. Informative tier makers are those for which the marker has been validated to test for the presence or absence of a given construct based on intensity. Partially informative tier makers were those for that could potentially be used to test for the presence or absence of a given construct based on intensity. Those markers which have not been tested were assigned the tier “Not tested”.

10) rsID.

11) Diagnostic. Name of the construct, substrain or strain group that the maker is diagnostic for. In all other cases is empty.

12) Diagnostic type. Substrain, strain group or construct.

13) Diagnostic information: Abbreviated name of the construct, name of substrain or list of substrains in which we observed the diagnostic allele. In all other cases is empty.

14) Partial diagnostic: 1, for diagnostic alleles that are not fixed. 0, in all other cases.

15) Diagnostic allele. Whether the reference or alternative allele is the diagnostic

16) Positive threshold. Threshold value to declare the presence of a given construct

17) Negative threshold. Threshold value to declare the absence of a given construct.

18) Uniqueness measured using Bowtie.

19) X chromosome markers used to determine the presence and number of X chromosomes. 1, chromosome X markers used in sex chromosome determination. 0, in all other cases.

20) Y chromosomes markers used to determine the presence of a Y chromosome. 1, chromosome Y markers used in sex chromosome determination. 0, in all other cases.

21) Flags. SPIKE, markers added in the final iteration of the array. Empty in all other cases.

22) Diagnostic Birth. The population where a diagnostic allele was first seen (E2, E3, E4 in the BxD, Pre-Cc, CC or Post-CC in the Collaborative Cross)

23) Diagnostic Fixation. The population where a diagnostic allele is inferred to be fixed (E3, CC or segregating)

**Supplementary Table 3.** List of inbred strains with consensus genotypes grouped into four classes: classical, wild-derived, CC and BXD.

**Supplementary Table 4.** Examples of the rules for consensus genotypes calls. *, denotes the diagnostic allele.

**Supplementary Table 5.** Consensus genotypes.

**Supplementary Table 6.** Aneuploid, mosaic and misclassified samples.

**Supplementary Table 7.** Number of samples with N and not N genotype calls in the autosomes and X chromosome of sample TL9348.

**Supplementary Table 8.** Construct probe design annotation

**Supplementary Figure 1.** Sex effect on normalized intensity for markers on chromosome X. The left figure represents 269 markers considered informative based on the lack of overlap between the distribution of intensities in males (blue) and females (red). The right represents 426 makers that are not considered sex informative.

**Supplementary Figure 2.** Alignments of validated construct markers. For each construct the file provides a short summary, the alignment of the working probes, the target DNA and protein sequences. The alignment of forward (black) and reverse (blue) probes is shown with the nucleotide used for “genotyping” (A) shown in red background for forward probes and in blue (T) for reverse probes. Mismatches are shown in purple background.

**Supplementary Figure 3.** Examples of normal and abnormal intensity distributions. Intensity distributions for six samples with low pd_stat and six samples with high pd_stat on the autosomes and chromosome X. Colored histogram bars are the intensity values distribution on the corresponding chromosome. Colored lines are the kernel density estimates for these data. Black lines are an attempt to fit the actual data to a normal curve.

**Supplementary Figure 4.** The distribution of pd_stat values in the 6,899 samples is shown on the y axis. The x axis shows the ks_stat for better contrast. Threshold determination for chromosomal sex using pd_stat. Samples in yellow were incorrectly identified as XO but are in fact XX (aka XX*, Supplementary Tables 1 and 4). Samples in green are from mouse species other than *Mus musculus*. Samples in blue are labeled Aneuploid by our algorithm. We manually established a threshold to capture all the misclassified samples and samples from other species.

**Supplementary Figure 5.** Chromosome Y duplications. Spatial distribution of normalized intensity at SNPs in the proximal end of the Y chromosome in C3H/He, DBA/1 and C57BL/6 samples. The range of intensities are shown in orange in cases where we had multiple samples with the duplication while samples with normal Y chromosome are shown in blue. Duplicated regions are shown in red and transition regions with uncertain copy number are shown in pink. The bottom of the figure shows the location of the MiniMUGA markers and genes.

**Supplementary Figure 6.** Intensities of all construct markers present in MiniMUGA. Markers are grouped according to construct. The color denotes whether the sample is deemed to be a negative control (blue), positive control (red), or experimental (dark brown) for the respective construct. Markers with asterisks were excluded in the construct analysis.

**Supplementary Figure 7.** Inbreeding thresholds. The figure shows in red the distribution of observed H calls in 385 samples representing 85 classical inbred strains. It also shows in blue the distribution of predicted number of H calls in 3,655 F1 hybrids using the consensus genotypes from 86 classical inbred strains. Tier 1 and 2 markers on the autosomes, X chromosomes and PAR were used. Thresholds for inbred, close to inbred and outbred are shown as vertical bars.

**Supplementary Figure 8.** MiniMUGA Background Analysis Report for the following four female cell lines: C2Cl2, GPG C3-Tag-T1-Luc, MLE12, and C57BL/6J.

**Supplementary Figure 9.** *De novo* X chromosome duplication. The range of intensities for females and males are shown in pink and blue, respectively. The sample with the duplication is shown as black line. Genotypes for the parental CC strains and the test sample are shown at the bottom as well as the first marker included in the duplication (asterisk) and the extent.

**Supplementary Figure 10.** Age and breeding history of four mouse samples from the B6.129-Nox4^tm1kkr^J congenic line maintained through breeding at UNC. Green triangles note the position of the generate allele. Red bars denote the ancestral allele for diagnostic SNPs fixed at E3. Pink bars denote ancestral alleles for diagnostic SNPs fixed at the start of the CC. Light blue bars denote diagnostic alleles at diagnostic SNPs fixed at E3. Lighter blue bars denote diagnostic alleles at diagnostic SNPs fixed at start of CC. Grey bars denote ancestral alleles at post-CC diagnostic SNPs. Dark blue bars denote diagnostic alleles at post-cc diagnostic SNPs. Split bars denote heterozygous SNPs in a sample.

## REFERENCES

1. Adams D. J., A. G. Doran, J. Lilue, and T. M. Keane, 2015 The Mouse Genomes Project: a repository of inbred laboratory mouse strain genomes. Mamm. Genome Off. J. Int. Mamm. Genome Soc. 26: 403–412. https://doi.org/10.1007/s00335-015-9579-6

2. Akeson E. C., L. R. Donahue, W. G. Beamer, K. L. Shultz, C. Ackert-Bicknell, et al., 2006 Chromosomal inversion discovered in C3H/HeJ mice. Genomics 87: 311–313. https://doi.org/10.1016/j.ygeno.2005.09.022

3. Arends D., S. Heise, S. Kärst, J. Trost, and G. A. Brockmann, 2016 Fine mapping a major obesity locus (jObes1) using a Berlin Fat Mouse × B6N advanced intercross population. Int. J. Obes. 2005 40: 1784–1788. https://doi.org/10.1038/ijo.2016.150

4. Ayabe S., K. Nakashima, and A. Yoshiki, 2019 Off- and on-target effects of genome editing in mouse embryos. J. Reprod. Dev. 65: 1–5. https://doi.org/10.1262/jrd.2018-128

5. Babbs R. K., J. A. Beierle, Q. T. Ruan, J. C. Kelliher, M. M. Chen, et al., 2019 Cyfip1 Haploinsufficiency Increases Compulsive-Like Behavior and Modulates Palatable Food Intake in Mice: Dependence on Cyfip2 Genetic Background, Parent-of Origin, and Sex. G3 Bethesda Md 9: 3009–3022. https://doi.org/10.1534/g3.119.400470

6. Carbonetto P., R. Cheng, J. P. Gyekis, C. C. Parker, D. A. Blizard, et al., 2014 Discovery and refinement of muscle weight QTLs in B6 × D2 advanced intercross mice. Physiol. Genomics 46: 571–582. https://doi.org/10.1152/physiolgenomics.00055.2014

7. Chesler E. J., D. M. Gatti, A. P. Morgan, M. Strobel, L. Trepanier, et al., 2016 Diversity Outbred Mice at 21: Maintaining Allelic Variation in the Face of Selection. G3 Bethesda Md 6: 3893–3902. https://doi.org/10.1534/g3.116.035527

8. Collaborative Cross Consortium, 2012 The genome architecture of the Collaborative Cross mouse genetic reference population. Genetics 190: 389–401. https://doi.org/10.1534/genetics.111.132639

9. Didion J. P., R. J. Buus, Z. Naghashfar, D. W. Threadgill, H. C. Morse, et al., 2014 SNP array profiling of mouse cell lines identifies their strains of origin and reveals cross-contamination and widespread aneuploidy. BMC Genomics 15: 847. https://doi.org/10.1186/1471-2164-15-847

10. Didion J. P., A. P. Morgan, L. Yadgary, T. A. Bell, R. C. McMullan, et al., 2016 R2d2 Drives Selfish Sweeps in the House Mouse. Mol. Biol. Evol. 33: 1381–1395. https://doi.org/10.1093/molbev/msw036

11. Dong Y., H. Li, L. Zhao, P. Koopman, F. Zhang, et al., 2019 Genome-Wide Off-Target Analysis in CRISPR-Cas9 Modified Mice and Their Offspring. G3 Bethesda Md 9: 3645–3651. https://doi.org/10.1534/g3.119.400503

12. Fragouli E., S. Alfarawati, D. D. Daphnis, N.-N. Goodall, A. Mania, et al., 2011 Cytogenetic analysis of human blastocysts with the use of FISH, CGH and aCGH: scientific data and technical evaluation. Hum. Reprod. Oxf. Engl. 26: 480–490. https://doi.org/10.1093/humrep/deq344

13. Heard E., and J. Turner, 2011 Function of the sex chromosomes in mammalian fertility. Cold Spring Harb. Perspect. Biol. 3: a002675. https://doi.org/10.1101/cshperspect.a002675

14. Johnson M., I. Zaretskaya, Y. Raytselis, Y. Merezhuk, S. McGinnis, et al., 2008 NCBI BLAST: a better web interface. Nucleic Acids Res. 36: W5–9. https://doi.org/10.1093/nar/gkn201

15. Johnson D. S., C. Cinnioglu, R. Ross, A. Filby, G. Gemelos, et al., 2010 Comprehensive analysis of karyotypic mosaicism between trophectoderm and inner cell mass. Mol. Hum. Reprod. 16: 944–949. https://doi.org/10.1093/molehr/gaq062

16. Keane T. M., L. Goodstadt, P. Danecek, M. A. White, K. Wong, et al., 2011 Mouse genomic variation and its effect on phenotypes and gene regulation. Nature 477: 289–294. https://doi.org/10.1038/nature10413

17. Kumar V., K. Kim, C. Joseph, S. Kourrich, S.-H. Yoo, et al., 2013 C57BL/6N mutation in cytoplasmic FMRP interacting protein 2 regulates cocaine response. Science 342: 1508– 1512. https://doi.org/10.1126/science.1245503

18. Le Gall J., M. Nizon, O. Pichon, J. Andrieux, S. Audebert-Bellanger, et al., 2017 Sex chromosome aneuploidies and copy-number variants: a further explanation for neurodevelopmental prognosis variability? Eur. J. Hum. Genet. EJHG 25: 930–934. https://doi.org/10.1038/ejhg.2017.93

19. McCoy R. C., 2017 Mosaicism in Preimplantation Human Embryos: When Chromosomal Abnormalities Are the Norm. Trends Genet. TIG 33: 448–463. https://doi.org/10.1016/j.tig.2017.04.001

20. Morgan A. P., C.-P. Fu, C.-Y. Kao, C. E. Welsh, J. P. Didion, et al., 2015 The Mouse Universal Genotyping Array: From Substrains to Subspecies. G3 Bethesda Md 6: 263–279. https://doi.org/10.1534/g3.115.022087

21. Morgan A. P., and F. Pardo-Manuel de Villena, 2017 Sequence and Structural Diversity of Mouse Y Chromosomes. Mol. Biol. Evol. 34: 3186–3204. https://doi.org/10.1093/molbev/msx250

22. Peirce J. L., L. Lu, J. Gu, L. M. Silver, and R. W. Williams, 2004 A new set of BXD recombinant inbred lines from advanced intercross populations in mice. BMC Genet. 5: 7. https://doi.org/10.1186/1471-2156-5-7

23. Rosshart S. P., B. G. Vassallo, D. Angeletti, D. S. Hutchinson, A. P. Morgan, et al., 2017 Wild Mouse Gut Microbiota Promotes Host Fitness and Improves Disease Resistance. Cell 171: 1015–1028.e13. https://doi.org/10.1016/j.cell.2017.09.016

24. Searle J. B., and R. M. Jones, 2002 Sex chromosome aneuploidy in wild small mammals. Cytogenet. Genome Res. 96: 239–243. https://doi.org/10.1159/000063017

25. Shorter J. R., F. Odet, D. L. Aylor, W. Pan, C.-Y. Kao, et al., 2017 Male Infertility Is Responsible for Nearly Half of the Extinction Observed in the Mouse Collaborative Cross. Genetics 206: 557–572. https://doi.org/10.1534/genetics.116.199596

26. Srivastava A., A. P. Morgan, M. L. Najarian, V. K. Sarsani, J. S. Sigmon, et al., 2017 Genomes of the Mouse Collaborative Cross. Genetics 206: 537–556. https://doi.org/10.1534/genetics.116.198838

27. Steemers F. J., W. Chang, G. Lee, D. L. Barker, R. Shen, et al., 2006 Whole-genome genotyping with the single-base extension assay. Nat. Methods 3: 31–33. https://doi.org/10.1038/nmeth842

28. Taylor B. A., H. J. Heiniger, and H. Meier, 1973 Genetic analysis of resistance to cadmium-induced testicular damage in mice. Proc. Soc. Exp. Biol. Med. Soc. Exp. Biol. Med. N. Y. N 143: 629–633. https://doi.org/10.3181/00379727-143-37380

29. Taylor B. A., C. Wnek, B. S. Kotlus, N. Roemer, T. MacTaggart, et al., 1999 Genotyping new BXD recombinant inbred mouse strains and comparison of BXD and consensus maps. Mamm. Genome Off. J. Int. Mamm. Genome Soc. 10: 335–348. https://doi.org/10.1007/s003359900998

30. Treger R. S., S. D. Pope, Y. Kong, M. Tokuyama, M. Taura, et al., 2019 The Lupus Susceptibility Locus Sgp3 Encodes the Suppressor of Endogenous Retrovirus Expression SNERV. Immunity 50: 334–347.e9. https://doi.org/10.1016/j.immuni.2018.12.022

31. Veale A. J., J. C. Russell, and C. M. King, 2018 The genomic ancestry, landscape genetics and invasion history of introduced mice in New Zealand. R. Soc. Open Sci. 5: 170879. https://doi.org/10.1098/rsos.170879

32. Yang H., Y. Ding, L. N. Hutchins, J. Szatkiewicz, T. A. Bell, et al., 2009 A customized and versatile high-density genotyping array for the mouse. Nat. Methods 6: 663–666. https://doi.org/10.1038/nmeth.1359

33. Yang H., J. R. Wang, J. P. Didion, R. J. Buus, T. A. Bell, et al., 2011 Subspecific origin and haplotype diversity in the laboratory mouse. Nat. Genet. 43: 648–655. https://doi.org/10.1038/ng.847

